# A new mechanism of regulation of LIM kinases, LIMK1 and LIMK2, modulates their activity on cofilin and actin filament remodelling

**DOI:** 10.64898/2026.06.23.733925

**Authors:** Elodie Villalonga-Rosso, Amandine Serrano, Cristine Gonçalves, Samia Aci-Sèche, Déborah Cassas, Célina Chalal, Bojan Žunar, Michel Doudeau, Christine Mosrin, Fabienne Godin, Pascal Bonnet, Hélène Bénédetti, Béatrice Vallée

## Abstract

LIM kinases, LIMK1 and LIMK2, play a crucial role in cytoskeleton dynamics. They are involved in many physiological processes but also in several pathologies such as cancer, neuronal diseases and neurofibromatosis. Although LIM kinases appear as promising therapeutic targets, they remain undruggable. A better understanding of their activity and regulation is thus required to better design efficient targeted therapies. Here, we have shown the impact of a single amino acid on LIMK activity on cofilin, their main substrate in actin filament remodelling. We demonstrated that Y632 and Y630, for LIMK1 and LIMK2 respectively, mediate LIMK dimerization, resulting in their transphosphorylation. This process seems to be a prerequisite for their canonical phosphorylation on their respective T508 and T505 residues within the activation loop. These Tyrosine are not phosphorylated, their aromatic nature is rather critical to ensure proper LIMK activity on cofilin. These results bring new insights into LIMK molecular features.

## Introduction

LIM kinases (LIMKs) are playing a crucial role in cell fate. Due to their role in cytoskeleton remodelling ^1–5^, they regulate many physiological processes such as the cell cycle ^6–11^, cell migration and motility ^12–17^, neuronal differentiation ^18–22^ and apoptosis ^23–26^. As a consequence, their implication in numerous pathologies has been shown: oncogenesis ^27–33^, viral infections ^34–38^, development of neurological and neurodegenerative diseases ^39–44^ and neurofibromatosis ^16,45–47^. As promising therapeutic targets, many small molecule inhibitors targeting their active site have been developed over the past 15 years. However, only one reached clinical stages phase I/IIa without published results ^48,49^. It is thus vital to have a better understanding of LIMK functions and regulations at the molecular level to further characterize these enzymes and develop successful therapeutic strategies for their inhibition.

LIM kinases are serine/threonine and tyrosine kinases. The LIM kinase family consists of only two members: LIMK1 and LIMK2, encoded by two different genes (ENSG00000106683, and ENSG00000182541, respectively) ^50^. These proteins are closely related, as they share 50% overall identity and 70% identity in their kinase domain ^50,51^. They are structured in two LIM (Lin-11, Isl-1 and Mec-3) domains at their N-terminus, each composed of two zinc fingers, a central PDZ domain, followed by an unstructured proline/serine (P/S) rich region, and a C-terminal kinase domain ^50^. In the literature, LIM kinases are mainly described for their role in cytoskeleton remodelling, notably through their activity on cofilin, a member of the Actin Depolymerizing Factor (ADF) family. When phosphorylated by LIM kinases on its serine 3 (S3), cofilin is inhibited, resulting in actin fiber polymerization and the formation of stress fibers ^1–3^. LIMKs also play a role in microtubule turnover, but the mechanism by which they are involved in this phenomenon has not yet been determined ^5,52^.

Canonically, LIM kinases act as downstream effectors of the members of the Rho GTPase family, including Rho, Rac and Cdc42, which modulate LIMK activity via their effectors, Rho-associated protein kinases (ROCK), myotonic dystrophy kinase-related Cdc42-binding kinases (MRCKα), and p21-activated kinases (PAK), PAK1, PAK2, and PAK4. LIMK1 and LIMK2 are activated by phosphorylation of their threonine 508 and threonine 505, respectively ^2,53^. However, it is now well-established that they have many more partners, regulators, substrates and are, in fact, at the very heart of a vast network of protein-protein interactions ^54^.

Our previous study of the isoforms of LIMK2 uncovered intriguing features on these proteins. Three isoforms of human LIMK2 are described in the literature: LIMK2a, LIMK2b, and LIMK2-1 ^55–58^. These three isoforms display different tissue-specific expressions, as well as different subcellular localizations, indicating that they may not have exactly the same functions ^58^. Furthermore, LIMK2 expression upon p53 activation after DNA damage is isoform-dependent: LIMK2b and LIMK2-1, but not LIMK2a, are upregulated, and lead to G2/M arrest via cofilin phosphorylation increase ^6^. LIMK2b and LIMK2-1 isoforms differ only from full-length LIMK2a at their extremities. The first LIM motif is truncated at the N-terminus of LIMK2b and LIMK2-1, whereas LIMK2-1 has an extra C-terminal domain identified as a Protein Phosphatase 1 inhibitory (PP1i) domain by sequence homology (**Figure 1**) ^55–57^. We have previously shown that contrary to LIMK2a and LIMK2b, LIMK2-1 has no kinase activity on cofilin, whereas it has a kinase activity on Myelin Basic Protein (MBP), a universal kinase substrate, and it is able to promote actin polymerization ^56^. In fact, we showed that LIMK2-1 acts as an inhibitor of the Protein Phosphatase 1, a protein that is known to dephosphorylate and activate cofilin^56^. Phosphatase inhibition activity of LIMK2-1 was recently thoroughly characterized ^59^. Thus, LIMK2-1 participates in the increase of the phosphorylated cofilin pool by preventing its dephosphorylation rather than by phosphorylating it ^56^. *Consequently, small molecule inhibitors targeting the LIMK active site might be inefficient towards LIMK2-1 and this isoform could constitute a bias in therapies targeting LIMKs*. These results strongly suggest that we are still missing a thorough understanding of the molecular determinants essential for LIMK activity on cofilin. Hence, it is necessary to further elucidate the molecular mechanisms required for the kinase activity of LIMKs on cofilin.

**Figure 1:**
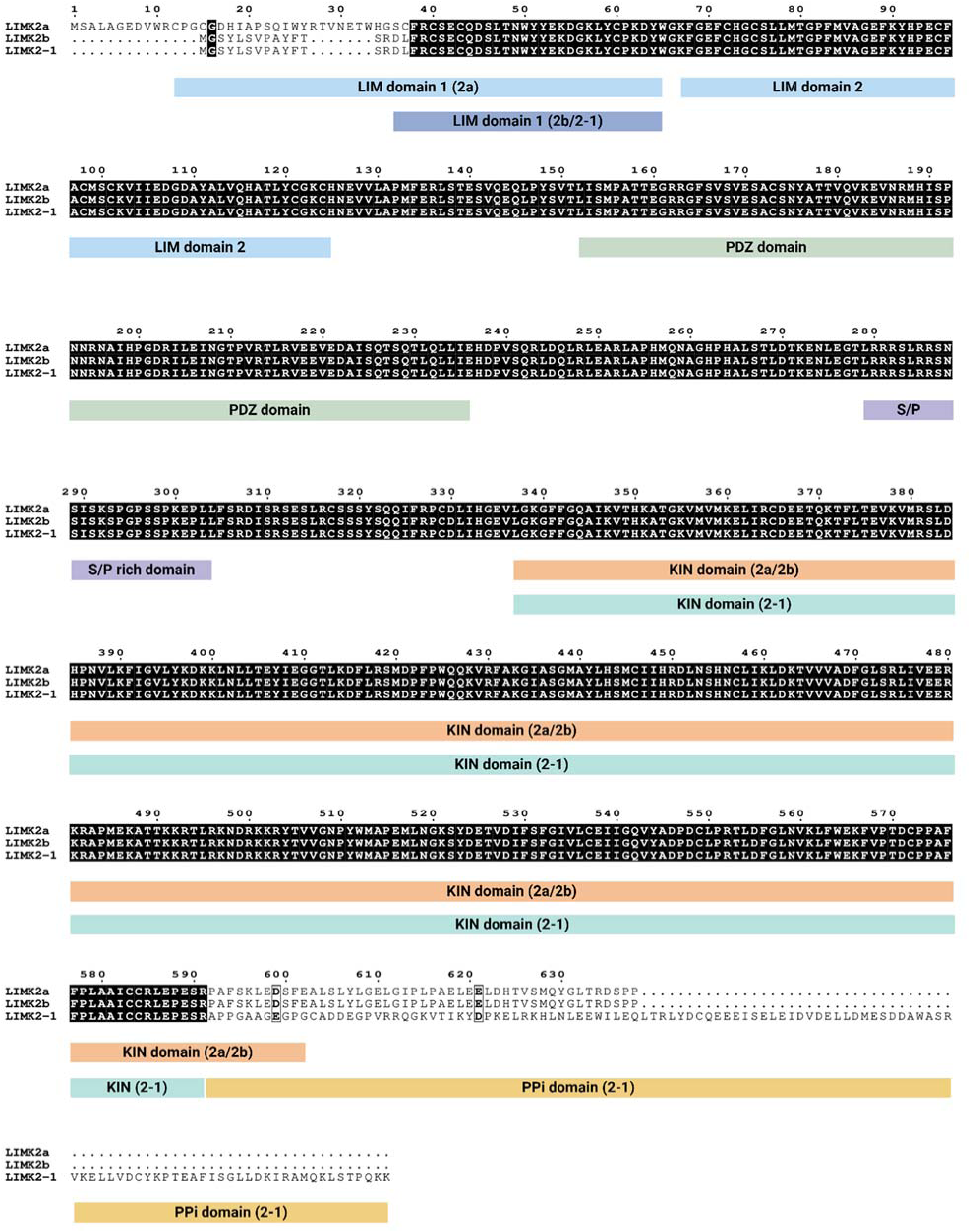
LIMK2 isoform alignment: the major differences between LIMK2a, 2b and 2-1 lie at their extremities. Sequence alignment and domain representation of the three isoforms of human LIMK2. LIMK2 isoforms are described in Entrez Gene: LIMK2-1 (NP_001026971.1), LIMK2a (NP_005560.1), LIMK2b (NP_057952.1). The various domains of LIMK2 are shown in color: LIM, PDZ, S/P rich (Serine Proline rich), KIN (Kinase), and PP1i (Protein Phosphatase 1 inhibitory). Conserved amino acids are written in white on a black background, conservative amino acids are written in bold in a frame.

In the present study, we have shown that the kinase domain of LIMK2 is not sufficient to phosphorylate cofilin, and that the very end of its C-terminal domain is required for cofilin phosphorylation. We have determined that a single amino acid, Y630 for LIMK2a and Y632 for LIMK1, is critical for their activity on cofilin. This amino acid mediates LIMK dimerization and transphosphorylation, which seems to be a prerequisite for LIMK2 and LIMK1 canonical activation by upstream kinases on T505 and T508, respectively. Our results unravelled an unexpected regulation of LIMK activation, and pave the way for a better understanding of their regulation.

## Materials and methods

### Materials

Anti-ROCK-1 (sc-17794) antibody was purchased from Santa Cruz Biotechnology Inc, anti-HA (11867423001) antibody from Roche Applied Science, anti-P-LIMK1 (T508)/LIMK2 (T505) (3841S), cofilin (5175S) and Phospho-cofilin (3313L) antibodies from Cell Signalling Technology. Anti-GFP antibody was purchased from Takara (632592), as well as TALON^®^ Metal Affinity Resin for 6xHis-tagged protein purification (635502). EZview™ Red anti-HA affinity gel (E6779) was purchased from Sigma-Aldrich Co, as well as anti-actin (A1978) antibody and cycloheximide (C-7898). GFP-Catcher beads were purchased from antibodies-online (ABIN5311508). Lipofectamine™ 2000 was purchased from Invitrogen (11668019). Recombinant MBP was purchased from Millipore (2243486).

### Expression and purification of cofilin 1

Plasmid carrying 6xHis-tag-*COFILIN 1* was transformed in Rosetta™(DE3)pLacI competent cells, according to the manufacturer instructions. Transformed colonies were selected on LB plates with ampicillin and chloramphenicol (100 µg/mL), used to generate 50 mL overnight cultures which were used to inoculate 2L of LB supplemented with ampicillin and chloramphenicol (100 µg/mL). Cultures were grown at 37 °C with shaking at 180 rpm until the OD600 nm reached 0.8. Expression of *cofilin-1* was then induced with 100 µM of isopropyl-β-D-thiogalactopyranoside (IPTG) at 16 °C overnight with shaking at 110 rpm. The induced cells were centrifuged at 4000 rpm for 10 minutes at 4 °C and resuspended in 30 ml of TBS buffer (20 mM Tris-HCl pH 8.0, 150 mM NaCl). The cells were then lysed using a sonicator in the presence of a cocktail of protease inhibitors (S8830, Sigma-Aldrich). Finally, cells were ultracentrifuged at 30,000 rpm for 30 minutes at 4 °C. Cofilin tagged with 6-histidine was purified by immobilized metal affinity chromatography using TALON Metal Affinity Resin. Non-specific proteins were eluted using low concentrations of imidazole (10-20 mM), and cofilin was eluted using 150 mM of imidazole. Fractions containing the fusion protein were then pooled and dialyzed overnight at 4 °C into TBS buffer at pH 8. A second dialysis was performed the following day for 4 hours. Cofilin was then concentrated using Amicon® Ultra-4 (UFC800324, Merck-Millipore) for further analysis.

### Plasmid construction

Plasmids were constructed by classical cloning using T4 DNA ligase upon digestion of amplified sequences of interest with Q5 high-fidelity DNA polymerase and restriction site insertion. Point mutations were created using Q5 site-directed mutagenesis kit. Plasmid DNA was isolated with NucleoSpin Plasmid Mini Kit and, when necessary, extracted from an agarose gel with NucleoSpin Gel and PCR Clean up Kit (Macherey-Nagel, Duren, Germany). All constructs were verified by restriction digest and Sanger sequencing. Primer synthesis and Sanger sequencing were conducted by Eurofins Genomics Europe (Luxembourg, Luxembourg). Oligonucleotides and plasmids used in this study are listed in supplementary data, **Table 1** and **Table 2**, respectively.

### Cell culture and transfection

HEK-293 (CRL-1573™) and HeLa cells (CRM-CCL-2™) were cultured under 5% CO_2_ at 37 °C in Dulbecco’s modified Eagle’s medium supplemented with 10% foetal calf serum. HEK-293 cells were transiently transfected with 10 μg of plasmid per 100-mm dish or 5 µg of plasmid per well of 6-well plates with Calcium Phosphate method. HeLa cells were transiently transfected with 0.8 µg of plasmid per well of 4-well plates with Lipofectamine 2000 according to manufacturer’s recommendations. Further experiments were conducted 48 hours after transfection.

### Cell lysates for protein detection

HEK-293 cells were lysed in 0.1 % Triton X-100 lysis buffer (50 mM Tris-HCl, pH 7.5, 100 mM NaCl, 5 mM EDTA, 50 mM NaF, 10 mM sodium pyrophosphate, 1 mM Na_3_VO_4_, 20 mM p-nitrophenyl phosphate, 20 mM β-glycerophosphate, 10 μg/mL aprotinin, 005 μg/mL okadaic acid, 1 μg/mL leupeptin, and 1 mM PMSF) on ice for 10 minutes. Upon 10-minute centrifugation at 10.000 x g, supernatants were collected and Laemmli buffer was added. Cell lysates were stored at −80 °C for further analysis by western blot.

### Immunoprecipitation

HEK-293 cells were transfected with expression plasmids as described above, and cultured for 48 hours. Cells were then lysed as described above. Supernatants were incubated for 2 hours at 4 °C either with anti-HA affinity gel for HA-LIMK2 constructs or with GFP-trap beads for YFP-LIMK2 constructs. Beads were then washed five times with lysis buffer, and then eluted with Laemmli sample buffer.

### Kinase assay

Immunoprecipitates bound to HA-beads or GFP-beads, as described above, were washed twice with lysis buffer and then three times with kinase buffer (50 mM HEPES-NaOH pH 7.5, 150 mM NaCl, 5 mM MgCl_2_, 5 mM MnCl_2_, 50 mM NaF, 1 mM Na_3_VO_4_, 20 mM β-glycerophosphate, 1 μg/mL leupeptin, and 1 mM PMSF). Immunoprecipitated beads were incubated for 20 minutes at 30 °C in 22 μl of kinase buffer containing 50 μM ATP, 5 μCi of γ[^32^P]ATP (3,000 Ci/mmol) and 2 μg of purified 6 x His-tag cofilin or 10 μg of commercial MBP. The reaction was terminated by heating 5 minutes at 90 °C in Laemmli sample buffer. Samples were then subjected to SDS-PAGE, and analysed by autoradiography.

### Cell staining

HeLa cells were fixed with 4% paraformaldelyde in PBS for 20 minutes and permeabilized with 0.5% Triton-X100 in PBS for 10 minutes at room temperature. After blocking with 0.1% Triton-X100, 2% BSA, 0.1% Azide in PBS for 10 minutes, cells were incubated with anti-HA antibody for 1 hour and subsequently with FITC-conjugated anti-rat IgG for 45 minutes. AlexaFluor568-conjugated phalloidin was added 20 minutes before the end of the secondary antibody incubation. The cells were washed three times with 0.1% Triton-X100 in PBS and nucleus was stained using Hoechst for 10 minutes. The cells were then washed three more times with 0.1% Triton-X100 in PBS, mounted on glass slides, and analyzed by fluorescence microscopy using Carl Zeiss Axio Observer Z7 (Carl Zeiss Co. Ltd., Iena, Germany). Images were acquired using ZEN v2.3 pro Zeiss software (Carl Zeiss Co. Ltd., Jena, Germany).

### Yeast growth assay

Cof^ts^ strain (cof1-5, ^60^) was transformed with different plasmids. These strains were grown overnight reaching stationary phase at 25 °C in selective SD-URA medium. The cultures were diluted at 0.1 OD, and grown in the same conditions until exponential phase (0.8 OD). A series of 5-fold dilutions were spotted onto selective agar plates containing 2% glucose or 2% galactose and grown either at 25 °C or 34 °C.

### Yeast cell extracts

Wild type BY4742 yeast strain was transformed with different plasmids. These strains were grown overnight reaching stationary phase at 30 °C in selective SD-URA medium. These subcultures were diluted and further grown until exponential phase (OD=0.8-1). Yeast cells were harvested by centrifugation, pellets were resuspended at 200 OD/ml in yeast lysis buffer (100 mM Tris-HCl pH 7.5, 150 mM NaCl, 5 mM EDTA, 1 mM PMSF, 1 µg/ml protease inhibitor mixture of Leupeptin, Pepstatin, Aprotinin). Cells were disrupted with glass beads. Cell debris and glass beads were then removed by centrifugation at 1.000 x g. Immunoprecipitation and kinase assay were performed as described above.

### Cycloheximide chase assay

HEK-293 cells were cultured and transfected with expression plasmids as described above. After 48 hours of transfection, cells were treated with 100 μg/mL of cycloheximide at the indicated chase times. At the end of the assay, cell samples were collected simultaneously, and lysed as described above. Cell lysates were stored at −80 °C before further analysis by western blot.

### Bioinformatics analysis

The alignment of protein sequences of the C-terminal extremity of LIMK2a in different species was obtained by performing Multiple Sequence Alignment (MSA) using the UniProt Align tool. The evolutionary rate of each amino acid position in the LIMK2 protein was generated using the ConSurf Data Base tool. Further analysis of the Y630A mutation impact on protein integrity were performed using PolyPhen-2 and Mutation t@ster prediction tools.

### Molecular modelling

The experimental structures of the kinase domain of the LIMKs proteins were retrieved from the RCSB PDB and superimposed by using the MatchMaker tool from the Chimera software. Among the 14 structures retrieved, PDB ID 3S95, 4TPT, 5HVJ, 5HVK, 5L6W, 5NXC, 5NXD, 7ATU, 7ATS, 7B8W, 7QHG, 8AAU, 8S3X, 8WSW, two representative structures of each conformational state of Y362 were chosen to carry out molecular dynamic simulations. The LIMK1 structure of PDB ID 3S95 represents the system with the Y632 in an “*out”* conformation while the LIMK1 structure of PDB ID 5NXC exhibits a residue Y632 in an “*in”* conformation. For both structures, the missing parts were completed by using the Modeller suite version 10.4 and the LIMK1 structure of PDB ID 5hvk.

#### System preparation

PROPKA version 3.5.1 ^61^ was used to check the protonation state of ionizable residue side-chains at a global pH value of 7.4. The AmberTools 24 ^62^ suite was employed to generate the topology and the coordinate file of each system. The protein force-field ff19SB parameters ^63^ were assigned. Then the system was solvated in a rectangular OPC water box with the side of the box being at least 12 Å away from any solute atom. Finally, counter ions were added to reach an ionic concentration of 0.15 M.

#### Molecular dynamics (MD) simulation

A three-cycle minimization was performed with 2000 steps each cycle, minimizing first the solvent, then the solute, and finally, the entire system. The SHAKE algorithm ^64^ was applied to constrain bonds involving hydrogen atoms, allowing a time increment of 2 fs.

Temperature regulation at 300 K was ensured through Langevin dynamics with a collision frequency of 2 ps^−1^. The long-range electrostatic interactions were computed using the Particle Mesh Ewald (PME) method beyond 10 Å distance. The system was slowly heated in canonical ensemble (NVT) from 0 to 300 K over a period of 20 ps, where a harmonic restraint on the solute (k=10 kcal·mol^−1^.Å^−1^) prevents the system from structural distortion. This was followed by a 30 ps equilibration step through which the harmonic restraints were gradually decreased from 10 kcal·mol^−1^·Å^−1^ to 1 kcal·mol^−1^·Å^−1^. The system was then equilibrated during a 1 ns MD simulation in the isobaric–isothermal ensemble (NPT) at 300 K and 1 atm. The pressure relaxation time was set to 1 ps. MD calculations were performed using the PMEMD.cuda module of the AMBER20 program ^61^. We performed 2 μs of MD production on each system and saved the coordinate every 1 ns.

### Statistics

All experiments were repeated at least three times. The data were analyzed and the figures were generated with Prism GraphPad 9 software (GraphPad, Inc., La Jolla, CA, USA). The results are presented as mean ± SEM of at least three experimental replicates. First, normal distribution of the data was assessed by the Shapiro-Wilk test. Statistical significance was then determined using one-way ANOVA followed by post-hoc tests using Tukey’s multiple comparisons test (*** P<0.001, ** P<0.01, * P<0.05).

## Results

### The C-terminal extremity of LIMK2 is required for its kinase activity on cofilin and its own phosphorylation

Major differences between LIMK2a/2b and LIMK2-1 lie at their C-terminal extremities (**Figure 1**). LIMK2-1 has a truncated kinase domain compared to LIMK2a (a few amino acids), followed by a PP1i domain. As LIMK2-1 has no kinase activity on cofilin, we thought that these missing amino acids may be required for its activity on cofilin.

We constructed two truncated mutants missing various lengths of the C-terminal extremity of the kinase domain of LIMK2a: LIMK2a-ΔCT591 and LIMK2a-ΔCT602 (**Figure 2A**). LIMK2a-ΔCT591 corresponds to the truncated kinase domain of LIMK2-1, whereas LIMK2a-ΔCT602 corresponds to the full-length kinase domain of LIMK2a, as described in sequence analysis databanks. We tested the ability of these mutants to phosphorylate cofilin in an *in vitro* assay. Plasmids carrying these different YFP-tagged mutants were transiently transfected in HEK-293 cells, which were then lysed. Lysates were immunoprecipitated with GFP-catcher beads. Immunoprecipitated beads were incubated with recombinant cofilin in the presence of γ[^32^P]ATP. [^32^P] incorporation into cofilin was assessed by autoradiography. It appeared that none of these two mutants was able to phosphorylate cofilin *in vitro* (**Figure 2B)**. We further characterized these mutants by testing their *in cellulo* activity on cofilin. Plasmids carrying the different HA-tagged mutants were transiently transfected in HEK-293 cells, which were then lysed. Western blot analysis of these lysates using an anti-LIMK2 antibody showed a very high expression of transfected forms of LIMK2 compared to endogenous LIMK2, suggesting that the contribution of endogenous LIMK2 is insignificant compared to transfected forms of LIMK2 (data not shown). Lysates were analyzed by western blot using anti-phospho-Ser3-cofilin antibody, as Ser3 of cofilin is phosphorylated by LIMK2. Neither of these two mutants was able to phosphorylate cofilin *in cellulo* (**Figure 2C**). Analysis of the 3D structure of the kinase domain of LIMK2a (PDB ID 4TPT) showed that the mutant LIMK2a-ΔCT602 is missing half of the penultimate α-helix of LIMK2a (**Figure 2A**). We then decided to construct two more mutants: LIMK2a-ΔCT609, bearing the full-length penultimate α-helix, and LIMK2a-ΔCT611, bearing two more amino acids, a glutamic acid and leucine, two residues that might be important for this helix conformation. We tested the activity of these two new mutants by our *in vitro* and *in cellulo* assays. Once more, neither of these two mutants was able to phosphorylate cofilin *in vitro* nor *in cellulo* (**Figure 2B and 2C**). These data suggest that the C-terminal part of LIMK2a (beyond amino acid 611) is required for LIMK2a kinase activity on cofilin both *in vitro* and *in cellulo*.

**Figure 2:**
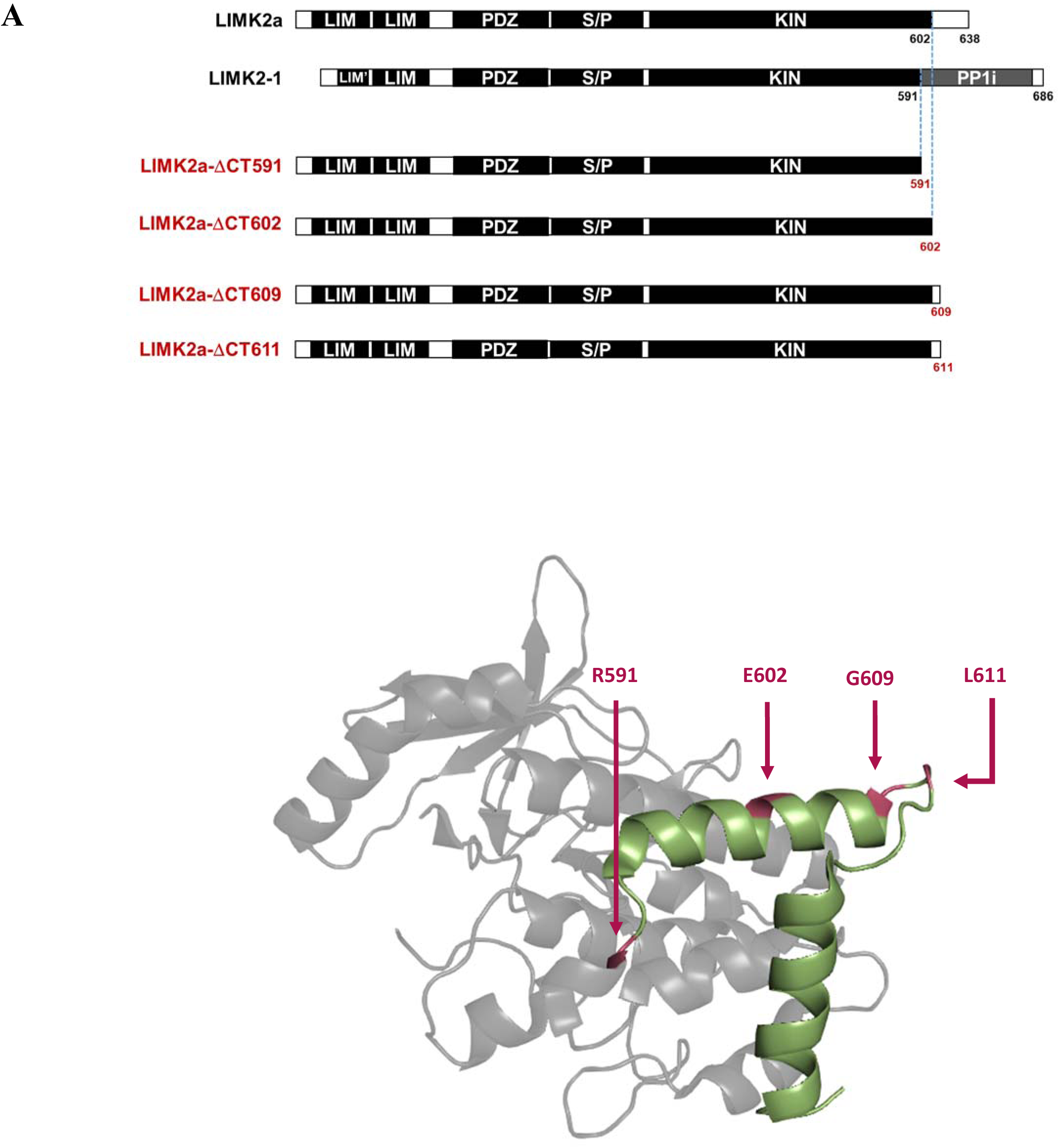
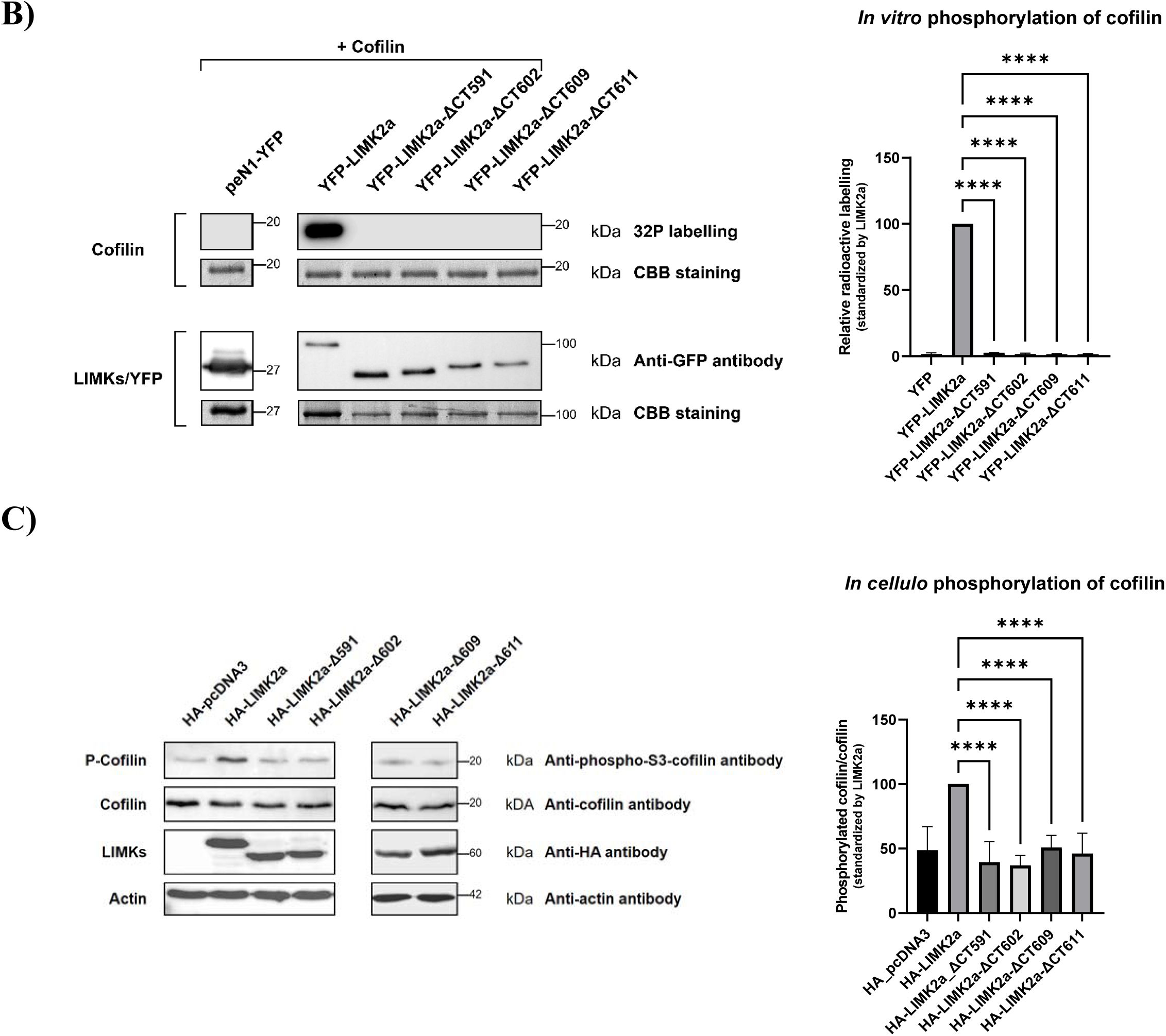
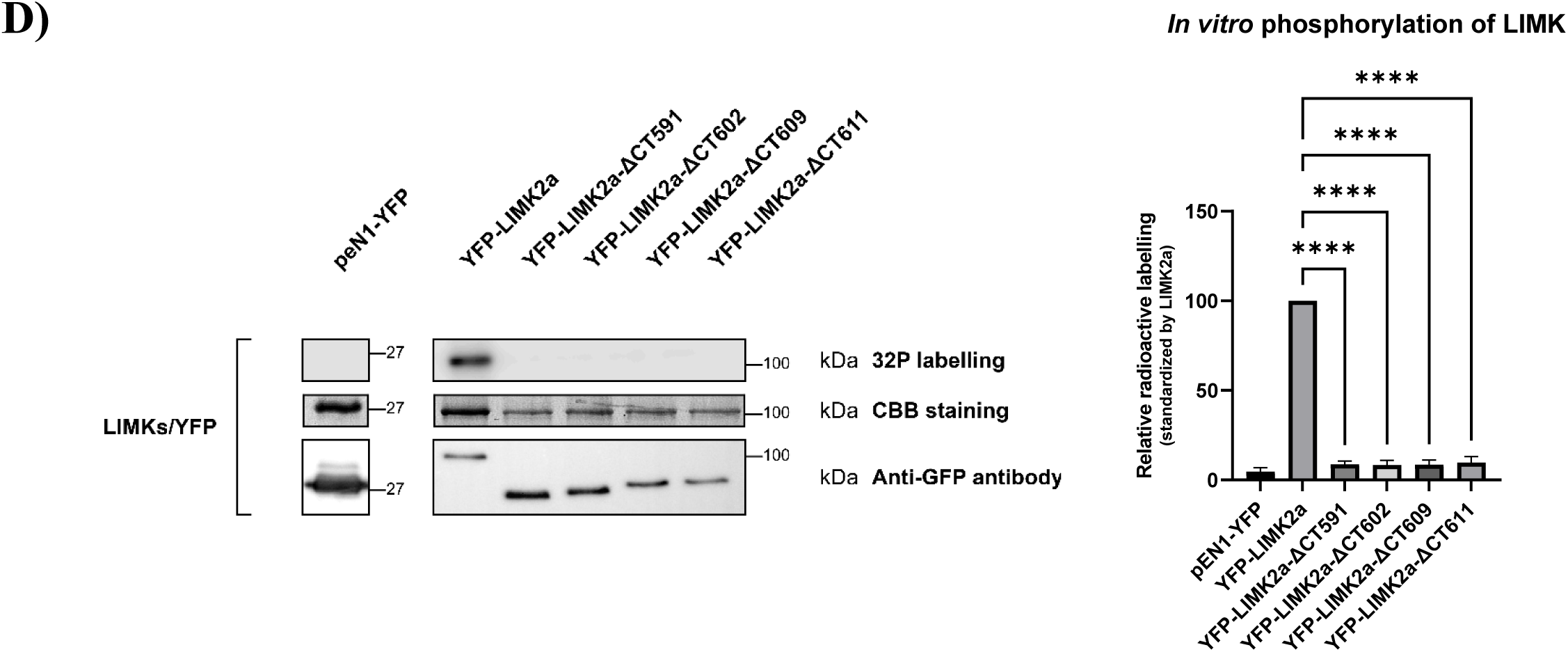
The C-terminal extremity of LIMK2 is required for its kinase activity on cofilin and its own phosphorylation. A) Schematic diagram and 3D structure of the C-terminal truncated LIMK2 constructions (PDB ID 4TPT). The two C-terminus helices are shown in green. B) *C-terminal truncated mutants of LIMK2a do not phosphorylate cofilin in vitro.* HEK-293 cells were transfected with plasmids allowing the overproduction of one of the four YFP-tagged LIMK2a truncated mutants (ΔCT591, ΔCT602, ΔCT609, ΔCT611) or YFP as a negative control. Anti-YFP immunoprecipitated proteins and recombinant 6xHis-cofilin were used in the kinase assay in the presence of γ[^32^P]ATP. The anti-YFP immunoprecipitates were also subjected to anti-GFP immunoblotting (10% acrylamide gel) and Coomassie blue staining (15% acrylamide gel). Quantification of the relative cofilin phosphorylation normalized by LIMK2a derivative CBB intensity is shown in the right graph. Wild-type transfected LIMK2a cell value was set at 100. C) *C-terminal truncated mutants of LIMK2a do not phosphorylate cofilin in cellulo.* HEK-293 cells were transfected with plasmids overproducing one of the four HA-tagged LIMK2a truncated mutants (ΔCT591, ΔCT602, ΔCT609, ΔCT611) or pcDNA3 as a negative control. Lysates were subjected to western blotting. Quantification of the ratio of phospho-cofilin versus cofilin normalized by LIMK2a derivatives is shown in the right graph. Wild-type transfected LIMK2a cell value was set at 100%. D) *C-terminal truncated mutants of LIMK2a are not phosphorylated in vitro.* HEK-293 cells were transfected with one of the four YFP-tagged LIMK2a truncated mutants (ΔCT591, ΔCT602, ΔCT609, ΔCT611) or YFP as a negative control. Anti-YFP immunoprecipitated proteins were used in the kinase assay in presence of γ[^32^P]ATP. The anti-YFP immunoprecipitates were also subjected to anti-GFP immunoblotting and Coomassie blue staining. Quantification of LIMK phosphorylation normalized by LIMK2a derivative CBB intensity is shown in the right graph. Wild-type transfected LIMK2a cell value was set at 100%.

Interestingly, while performing the kinase assay on cofilin, we observed that the four truncated mutants were no longer phosphorylated in our *in vitro* assay contrary to wild-type LIMK2a (**Figure 2D**). These last data suggest that one or several residues at the very end of the C-terminal extremity (between amino acids 612 and 638) of LIMK2a seem required for LIMK2a phosphorylation.

### LIMK2a-Y630A mutant does not phosphorylate cofilin

We then decided to determine which amino acid(s) of the C-terminal part of LIMK2a contribute to LIMK2 phosphorylation. At the very C-terminal extremity of LIMK2a, five amino acids (T625, S627, Y630, T633 and S636) are potential sites of phosphorylation (**Figure 3A**, marked by a red asterisk). By site-directed mutagenesis, we mutated them one by one into alanine residues, and tested the kinase activity of each of these mutants on cofilin.

**Figure 3:**
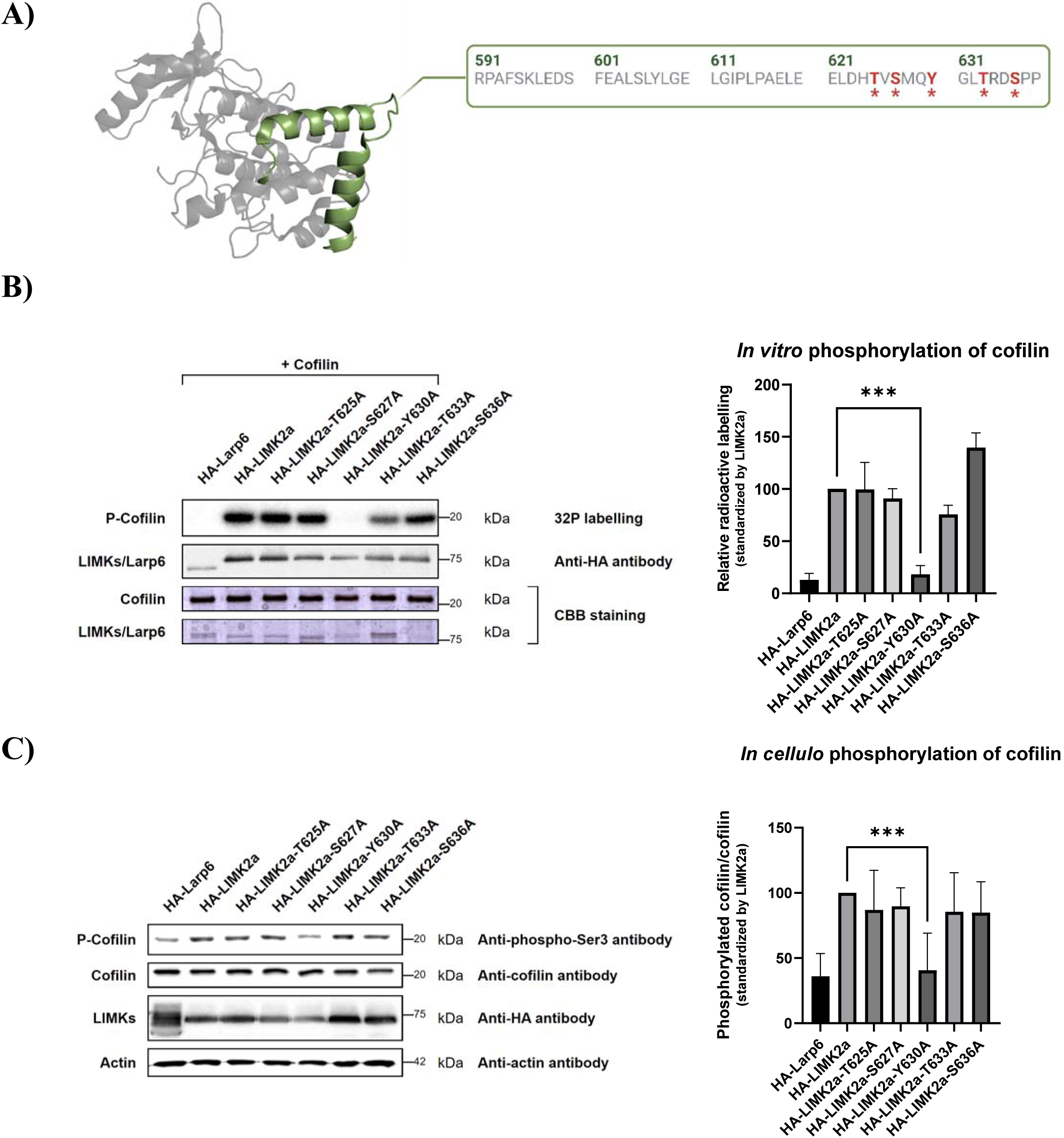
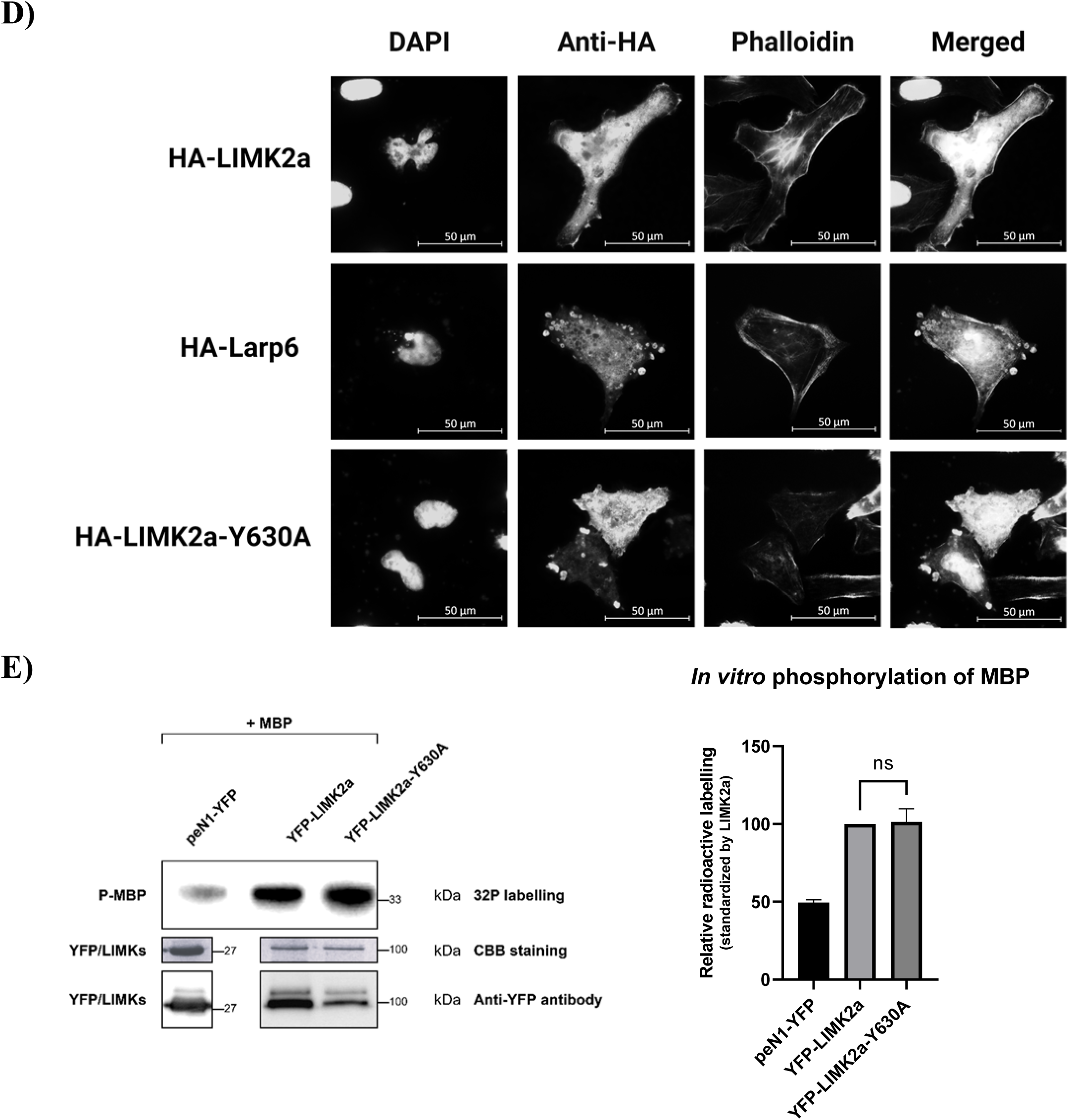
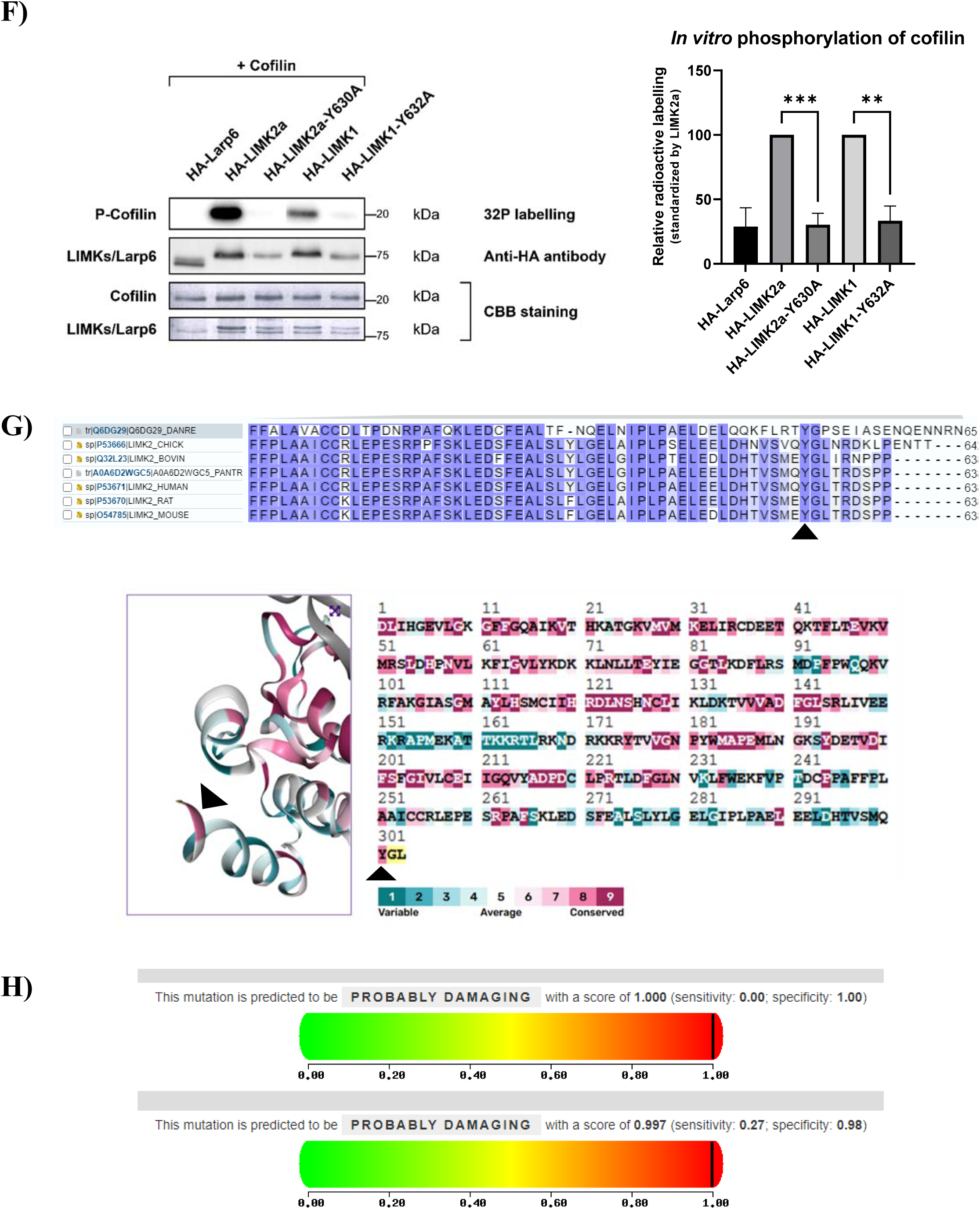
LIMK2a-Y630A mutant does not phosphorylate cofilin. A) *3D structure of the kinase domain of LIMK2a (PDB ID 4TPT) and sequence of the C-terminal extremity of LIMK2a.* Potential phosphorylated amino-acids corresponding to site-directed mutants are indicated by a red asterisk: T625, S627, T630, T633 and S636. B) *LIMK2a-Y630A does not phosphorylate cofilin in vitro.* HEK-293 cells were transfected with plasmids allowing the overproduction of indicated proteins. Larp6, an unrelated HA-tagged protein, was used as a negative control. Anti-HA immunoprecipitated proteins were used in the kinase assay. The anti-HA immunoprecipitates were also subjected to anti-HA immunoblotting and Coomassie blue staining. Quantification of cofilin phosphorylation as depicted in Figure 2B is shown in the right graph. C) *LIMK2a-Y630A does not phosphorylate cofilin in cellulo.* HEK-293 cells were transfected with plasmids allowing the overproduction of indicated proteins. Larp6 was used as a negative control. Lysates were subjected to western blotting. Quantification of cofilin phosphorylation as depicted in Figure 2C is shown in the right graph. D) *LIMK2a-Y630A is unable to induce stress fibers.* HeLa cells were transfected with plasmids overproducing HA-tagged WT LIMK2a, the mutant LIMK2a-Y630A or HA-tagged Larp6 as a negative control. Cells were fixed and actin cytoskeleton was stained using phalloidin. Anti-HA antibody labelling was used to show transfected cells. The scale bar represents 50 µm. E) *LIMK2a-Y630A is able to phosphorylate MBP in vitro.* HEK-293 cells were transfected with plasmids allowing the overproduction of YFP-tagged LIMK2a or LIMK2a-Y630A. Plasmid peN1-YFP was used as a negative control. Anti-YFP immunoprecipitated proteins and recombinant MBP were used in the kinase assay in presence of γ[^32^P]ATP. The anti-YFP immunoprecipitates were also subjected to anti-GFP immunoblotting and Coomassie blue staining. Quantification of cofilin phosphorylation as depicted in Figure 2B is shown in the right graph. F) *LIMK1-Y632A does not phosphorylate cofilin in vitro.* HEK-293 cells were transfected with plasmids allowing the overproduction of indicated proteins. HA-tagged protein Larp6 was used as a negative control. Anti-HA immunoprecipitated proteins were used in the kinase assay. The anti-HA immunoprecipitates were also subjected to anti-HA immunoblotting and Coomassie blue staining. Quantification of cofilin phosphorylation as depicted in Figure 2B is shown in the right graph. Wild-type LIMK2a or wild-type LIMK1 transfected cell values were set at 100%. G) *Tyrosine 630 is highly conserved in LIMK2a*. Top panel: Alignments of protein sequences of the C-terminal extremity of LIMK2a in different species: Danio rerio (Q6DG29), Gallus gallus domesticus (P53666), Bos taurus (Q32L23), Pan troglodytes (A0A6D2WGC5), Homo sapiens (P53671), Rattus norvegicus (P53670), Mus musculus (O54785). Tyrosine 630 is indicated by a black arrowhead. Bottom panel: Evolutionary rate of each amino acid position in LIMK2 protein using ConSurf Data Base on PDB 4TPT. Y630 is indicated by black arrowheads. H) *Analysis of Y630A mutation by bioinformatics tools predicts highly damaged protein.* Predictive effects of the mutation were evaluated using two different bioinformatics tools: Polyphen2 (upper panel) and MutationTaster (lower panel).

*In vitro* as well as *in cellulo* assays showed that LIMK2a-Y630A was unable to phosphorylate cofilin whereas the other mutants phosphorylated cofilin at the same level as wild-type LIMK2a (**Figures 3B and 3C**). As LIMK2 mutants were differently expressed, all the quantifications of cofilin phosphorylation were standardized by LIMK2 expression. The same quantification process was applied for all of the following experiments.

We then looked at cytoskeleton remodelling induced by LIMK2a by immunofluorescence, by labelling actin filaments with phalloidin. Indeed, LIMK2 overexpression is known to induce actin polymerization leading to stress fiber formation. HeLa cells were transfected with a plasmid allowing the expression of either wild-type LIMK2a, LIMK2a-Y630A or the unrelated protein Larp6. Contrary to wild-type LIMK2a, LIMK2a-Y630A was unable to induce stress fibers, suggesting that this mutant is inactive on actin filament polymerization (**Figure 3D**). This result is in good agreement with our *in vitro* and *in cellulo* results.

We then tested if LIMK2a-Y630A loss of kinase activity was cofilin-specific or if it was more general towards other substrates, thus suggesting a dysfunctional kinase domain. We used MBP, Myelin Basic Protein, a protein known to be a broad substrate for many kinases, and tested if LIMK2a-Y630A could phosphorylate it in our *in vitro* assay (**Figure 3E**). LIMK2a-Y630A did phosphorylate MBP, suggesting that its kinase domain is functional. LIMK2a-Y630A’s loss of kinase activity seems to be specific of its cofilin substrate.

We then checked if the role of this tyrosine 630 was also conserved in LIMK1. Y630 of LIMK2 corresponds to Y632 in LIMK1. We constructed the LIMK1-Y632A mutant and tested its activity on cofilin in *our vitro* assay. LIMK1-Y632A was unable to phosphorylate cofilin (**Figure 3F**). These results confirm that the role of this tyrosine is conserved in both LIMK1 and LIMK2.

Along this line, we analyzed the conservation of this tyrosine among evolution. BLAST alignments revealed that this tyrosine is extremely conserved among species (**Figure 3G**, top panel). ConSurf analysis was then performed to assess the evolutionary rate of each amino acid position in the LIMK2a sequence. This highlights the conservation of this tyrosine among evolution and highly suggests the functional importance of this residue (**Figure 3G**, bottom panel). We also wanted to assess the consequence of the mutation of tyrosine 630 in protein function using Polyphen and Mutation Taster tools. Both analyses revealed that mutation of Y630 is deleterious for LIMK2a suggesting a strong impact of this amino acid on LIMK2 structure and function (**Figure 3H**).

### LIMK2a-Y630A mutant downstream and upstream partners

As LIMK2a-Y630A is unable to phosphorylate its substrate cofilin, we wondered if it was still able to interact with it. By co-immunoprecipitation experiments, we showed that transfected YFP-LIMK2a-Y630A interacted with endogenous cofilin (**Figure 4A**). Lack of kinase activity of LIMK2a-Y630A on cofilin is thus not due to a loss of interaction between this mutant and cofilin.

**Figure 4:**
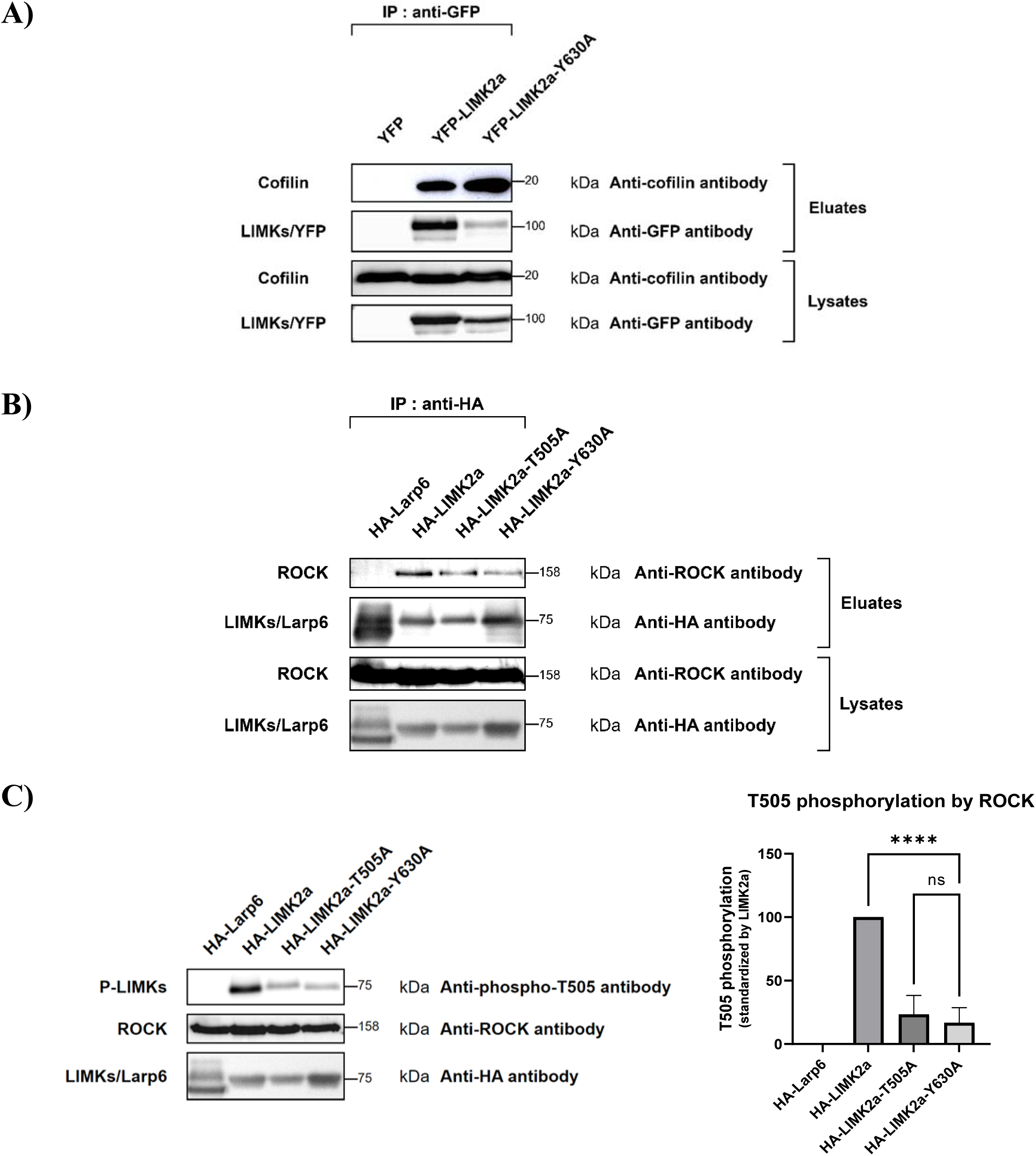
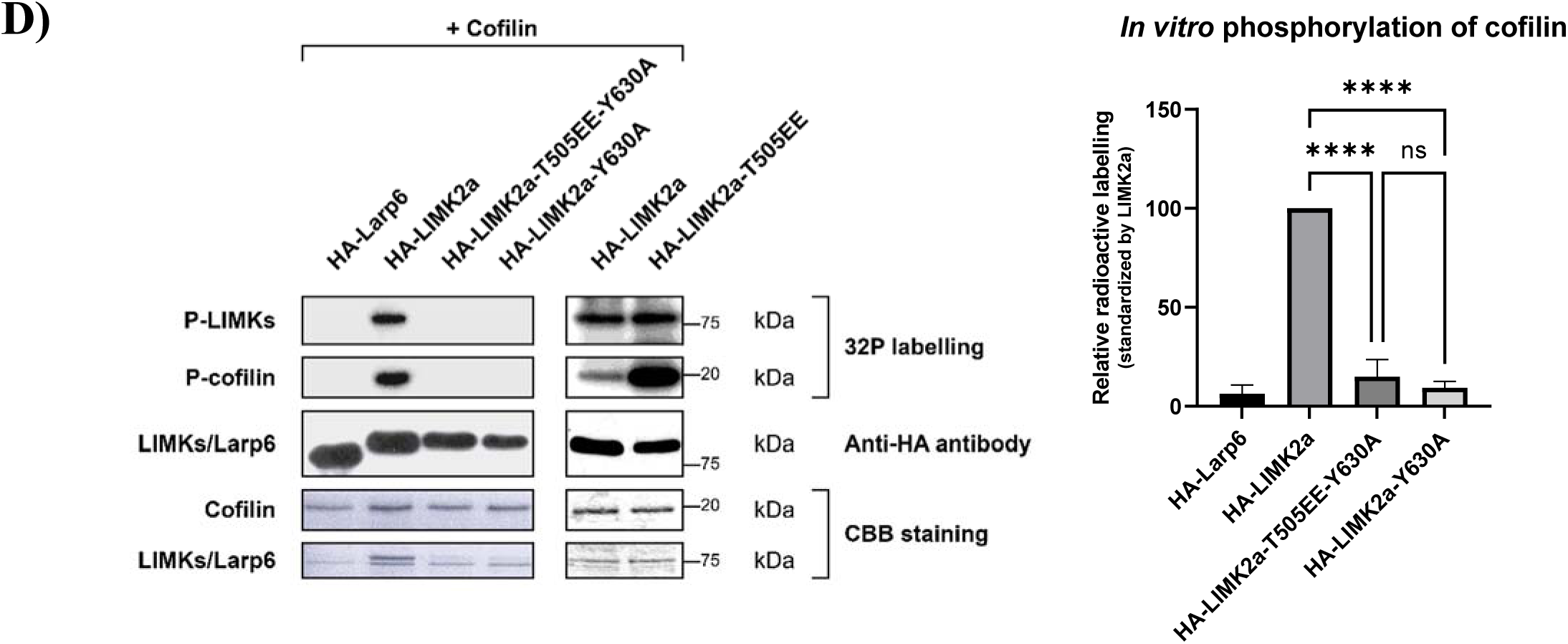
LIMK2a-Y630A mutant downstream and upstream partners and activation on its T505. A) *LIMK2a-Y630A interacts with endogenous cofilin.* HEK-293 cells overproduced indicated proteins and lysed. YFP-peN1 was used as a negative control. Lysates were immunoprecipitated with GFP-catcher beads. Lysates and anti-YFP immunoprecipitates were subjected to western blotting. B) *LIMK2a-Y630A is able to interact with the upstream kinase ROCK.* HEK-293 cells were co-transfected with plasmids allowing the overproduction of indicated proteins and cMyc-tagged ROCK and lysed. Larp6 was used as a negative control. Lysates were immunoprecipitated with anti-HA beads. Lysates and anti-HA immunoprecipitates were subjected to western blotting. C) *LIMK2a-Y630A is not phosphorylated by ROCK on T505.* HEK-293 cells were co-transfected with plasmids allowing the overproduction of indicated proteins and cMyc-tagged ROCK and lysed. Larp6 was used as a negative control. Lysates were subjected to western blotting. Quantification of T505 phosphorylation of LIMK2a normalized by LIMK2a derivatives is shown in the right graph. Wild-type LIMK2a transfected cell value was set at 100%. D) *LIMK2a-T505EE-Y630A does not phosphorylate cofilin in vitro*. HEK-293 cells overexpressed indicated proteins. Anti-HA immunoprecipitated proteins were used in the kinase assay. The anti-HA immunoprecipitates were also subjected to anti-HA immunoblotting and Coomassie blue staining. Quantification of cofilin phosphorylation as depicted in Figure 2B is shown in the right graph.

ROCK is one of the upstream kinases activating LIMK2 by phosphorylation on its threonine 505. We assessed the interaction between LIMK2a and ROCK. By co-immunoprecipitation experiments on transfected ROCK and LIMKs, we showed that LIMK2a-Y630A interacted with ROCK (**Figure 4B**). Then, we checked if LIMK2a-Y630A was still activated by ROCK via the phosphorylation of its T505 by western-blot analysis using an anti-phospho-T505-LIMK specific antibody on cell lysates co-transfected by plasmids overproducing LIMKs and ROCK. LIMK2a-Y630A phosphorylation on its T505 was drastically decreased compared to wild-type LIMK2, and appeared to be at the same level as the LIMK2-T505A mutant which, because it cannot be phosphorylated on its T505, serves as a negative control for this antibody, which has a bit of background noise. It then appeared that ROCK did not phosphorylate LIMK2a-Y630A on its T505 thus impairing its canonical activation (**Figure 4C**).

As LIMK2a-Y630A lacks canonical phosphorylation/activation on its T505, this may explain its loss of activity on cofilin. We then decided to investigate if we could bypass the impact of the Y630A mutation on cofilin phosphorylation by taking advantage of the constitutively active mutant LIMK2a-T505EE previously described in the literature ^53^. We constructed a double LIMK2 mutant bearing both T505EE and Y630A mutations. We performed our *in vitro* assay with this new mutant and compared its activity on cofilin with wild-type LIMK2a, LIMK2a-T505EE and LIMK2a-Y630A mutants. Surprisingly, this double mutant was not able to phosphorylate cofilin (**Figure 4D)**. Our results suggest that LIMK2 activation by phosphorylation of its T505 in its activation loop is not sufficient for its kinase activity on cofilin. Y630 seems to trigger a process crucial for LIMK2 kinase activity on cofilin which seems independent of its canonical phosphorylation on T505. Furthermore, in this experiment, LIMK2a phosphorylation was also monitored. No phosphorylation of LIMK2a-Y630A was detected. This may be explained by the lack of phosphorylation on its T505. However, LIMK2a-T505EE was phosphorylated in this assay, suggesting that another amino acid beside T505 is phosphorylated. Interestingly, no phosphorylation of LIMK2a-Y630A-T505EE was detected either (**Figure 4D**).

### How does tyrosine 630 affect LIMK2 kinase activity on cofilin and LIMK2 phosphorylation?

As Y630A mutation led to a complete loss of LIMK2 as well as LIMK2-T505EE phosphorylation in our *in vitro* assays (**Figure 4D**), we hypothesized that this tyrosine may be phosphorylated and that this phosphorylation might be a prerequisite for T505 phosphorylation. This would explain the loss of phosphorylation of LIMK2a-Y630A on its T505 (Figure 4C). Both phosphorylation of Y630 and T505 would then be required for optimal activity of LIMK2 on cofilin as the mutant LIMK2a-Y630A-T505EE does not phosphorylate cofilin.

First, we mutated tyrosine 630 into another phosphorylatable amino acid, serine (S). We performed our *in vitro* assay of cofilin phosphorylation with this new mutant. LIMK2a-Y630S was unable to phosphorylate cofilin (**Figure 5A**), and was not phosphorylated in our *in vitro* assay (**Figure 5B**).

**Figure 5:**
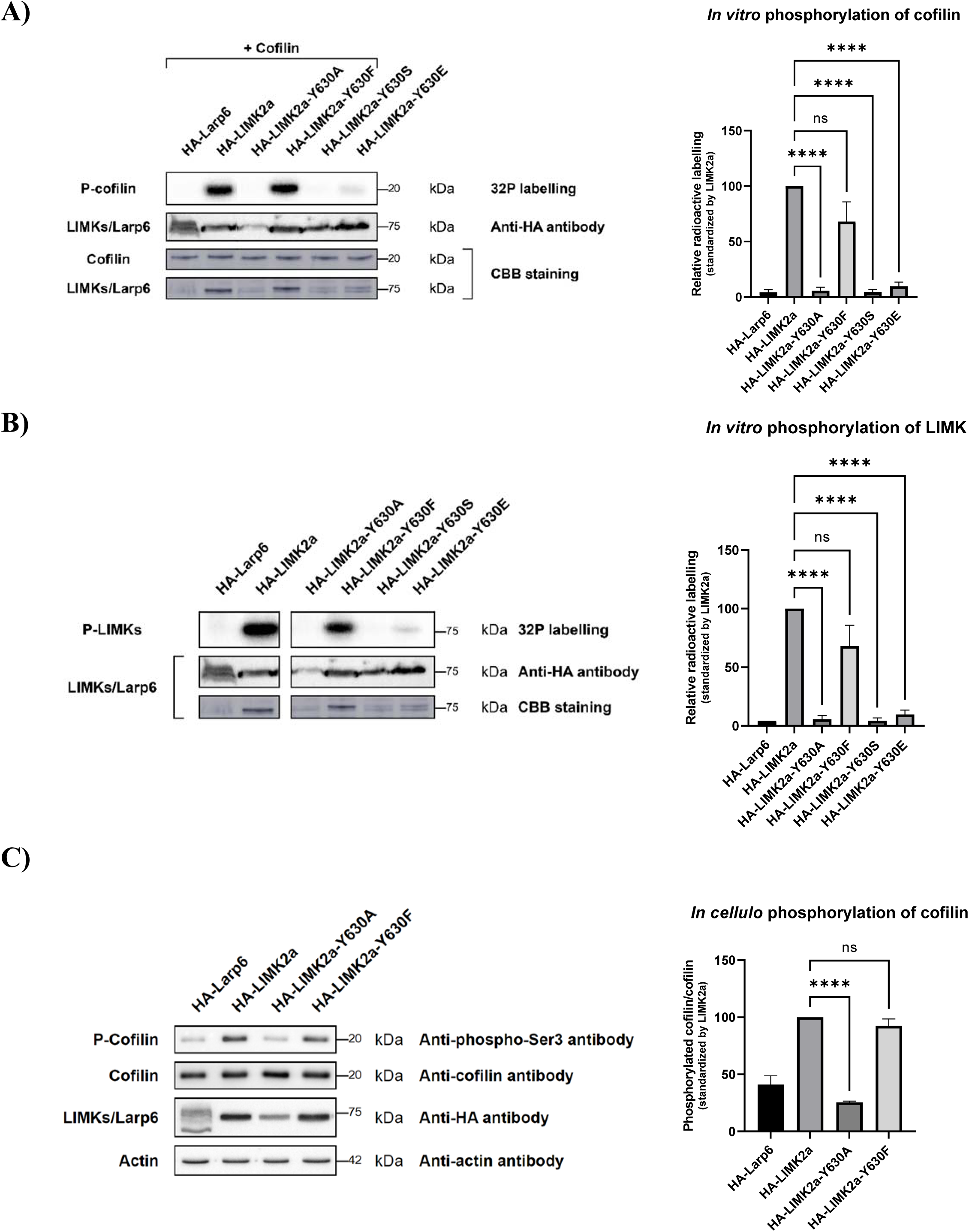
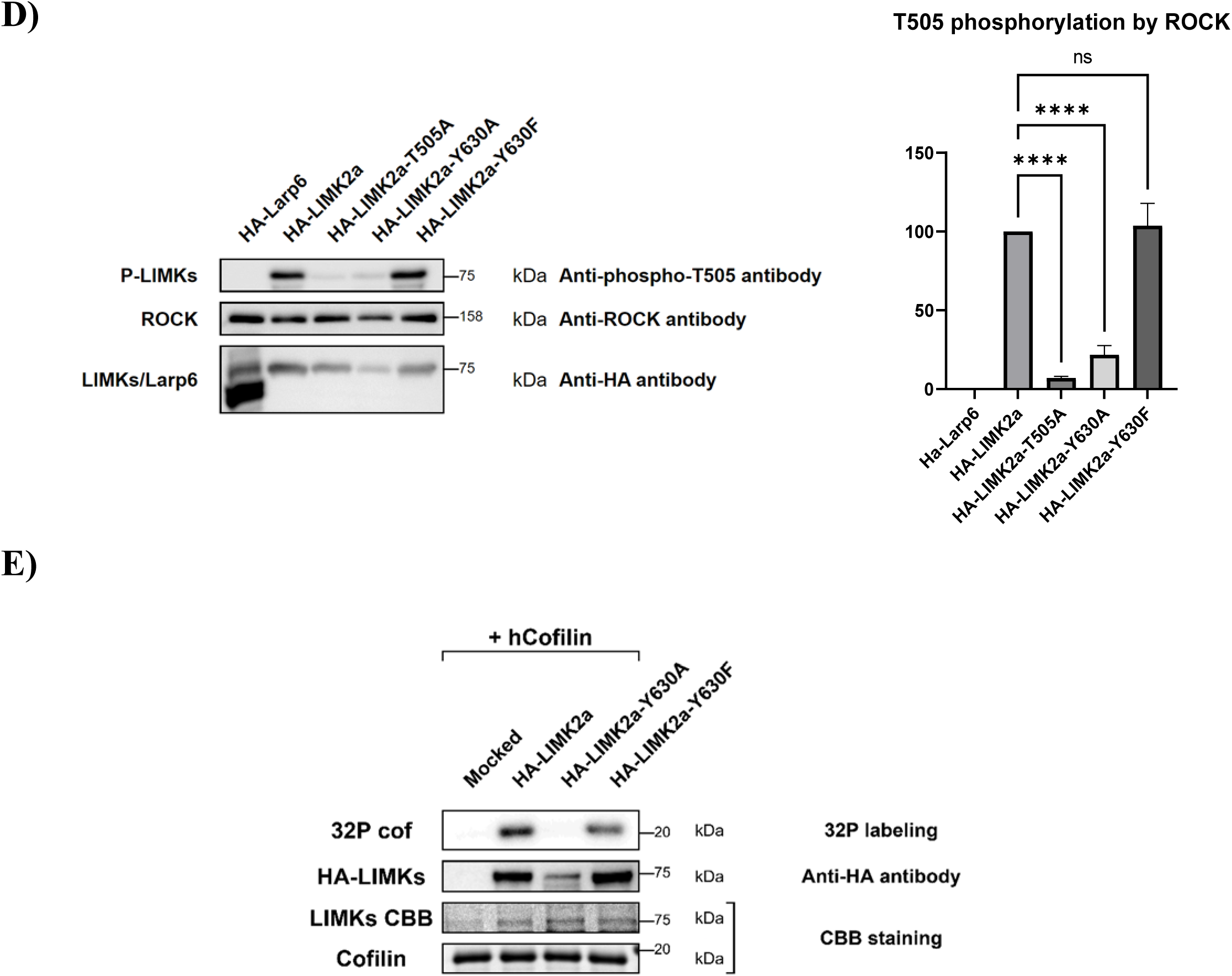
How is tyrosine 630 affecting LIMK2 kinase activity on cofilin? A) *LIMK2a-Y630F restores cofilin phosphorylation in vitro.* HEK-293 cells overproduced indicated proteins. Anti-HA immunoprecipitated proteins were used in the kinase assay. The anti-HA immunoprecipitates were also subjected to anti-HA immunoblotting and Coomassie blue staining. Quantification of cofilin phosphorylation as depicted in Figure 2B is shown in the right graph. B) *LIMK2a-Y630F is phosphorylated in vitro.* HEK-293 cells overproduced indicated proteins. Anti-HA immunoprecipitated proteins were used in the kinase assay. The anti-HA immunoprecipitates were also subjected to anti-HA immunoblotting and Coomassie blue staining. Quantification of LIMK2a phosphorylation as depicted in Figure 2D is shown in the right graph. C) *LIMK2a-Y630F phosphorylates cofilin in cellulo.* HEK-293 cells overproduced indicated proteins. Lysates were subjected to western blotting. Quantification of phospho-cofilin as depicted in Figure 2C is shown in the right graph. D) *LIMK2a-Y630F is phosphorylated by ROCK on its T505.* HEK-293 cells overproduced with indicated proteins and c-Myc-tagged ROCK. Lysates were subjected to western blotting. Quantification of T505 phosphorylation of LIMK2a as depicted in Figure 4C is shown in the right graph. E) *LIMK2a-Y630A overexpressed in yeast does not phosphorylate cofilin*. BY4741 yeast strain was transformed with plasmids allowing the overproduction of the different LIMK2 mutants. Cells were lysed and subjected to anti-HA immunoprecipitation. *In vitro* phosphorylation assay was performed on these immunoprecipitated beads.

We then decided to mutate Y630 into the phosphomimetic amino acid glutamic acid (E). The phosphomimetic amino acid glutamic acid (E) is commonly used to mimic phospho-serine or phospho-threonine, but it has also been shown to efficiently mimic phospho-tyrosine ^65^. We then performed our *in vitro* assay with this mutant. LIMK2a-Y630E mutant was unable to phosphorylate cofilin (**Figure 5A**) and was not phosphorylated on its own (**Figure 5B**).

Our results suggested that our Y630 phosphorylation hypothesis may not be accurate. Instead, we hypothesized that Y630 may alternatively play a role related to its structure, as it contains an aromatic residue. We decided to mutate Y630 into phenylalanine (F), a closely related amino acid from a structural point of view. We performed our *in vitro* cofilin phosphorylation assay with this new mutant. Interestingly, this mutant was able to phosphorylate cofilin at the same level as wild-type LIMK2a (**Figure 5A**). We also checked *in cellulo* phosphorylation of cofilin mediated by this mutant. It also restored wild-type level of phospho-cofilin (**Figure 5C**). Furthermore, LIMK2a-Y630F mutant was phosphorylated in our *in vitro* assay (**Figure 5B**) suggesting that Y630 is not a phosphorylation site. Furthermore, ROCK phosphorylation on T505 was also fully restored for this mutant (**Figure 5D**).

### Contrary to LIMK2a and LIMK2a-Y630F, LIMK2-Y630A does not phosphorylate cofilin in yeast

To rule out the potential involvement of partners copurified with LIMK2a in the mammalian system, we overexpressed the different mutants of LIMK2a in an orthologous system, *i.e.* a wild type strain of *Saccharomyces cerevisiae.* LIMKs are not present in the yeast *S. cerevisiae,* appearing later in the evolution. We then assessed yeast purified LIMK2a mutant’s ability to phosphorylate cofilin by performing our *in vitro* assay. In these conditions, LIMK2a as well as LIMK2a-Y630F phosphorylated cofilin, whereas LIMK2a-Y630A did not (**Figure 5E**).

### LIMK2 phosphorylation occurs by a transphosphorylation triggered by its dimerization

Our *in vitro* results suggest that Y630 triggers one or several phosphorylation(s) of LIMK2 apart from T505 and itself, which are essential for its kinase activity on cofilin (**Figures 4D and 5B**). We assessed how the phosphorylation process mediated by Y630 may occur. Three hypotheses were explored: i) the phosphorylation could be triggered by another kinase, ii) it could result from a transphosphorylation, or iii) from an autophosphorylation. Indeed, LIM kinases are known for their ability to dimerize and to transphosphorylate as well as to autophosphorylate ^66^.

To decipher between these three different possibilities, we took advantage of two mutants well-described in the literature: LIMK2a-D451N and LIMK2a-P386E. LIMK2a-D451N is a kinase-dead mutant of LIMK2a, while P386E mutation corresponds to LIMK1-P394E mutation which prevents LIMK1 dimerization and transphosphorylation ^67^. We performed our *in vitro* assay with these two mutants. As expected, LIMK2a-D451N was unable to phosphorylate cofilin. Moreover, it was not phosphorylated on its own (**Figure 6A**). This result shows that LIMK2 kinase activity is required for its own phosphorylation ruling out the implication of another kinase. LIMK2a-P386E did not phosphorylate cofilin and was not phosphorylated (**Figure 6A**). These results show that LIMK2a phosphorylation is extinguished when its dimerization is disrupted. Altogether, our results indicate that LIMK phosphorylation is neither the result of a phosphorylation by another kinase, nor the result of an autophosphorylation, but is rather the result of a transphosphorylation due to its dimerization, a process required for LIMK activity on cofilin.

**Figure 6:**
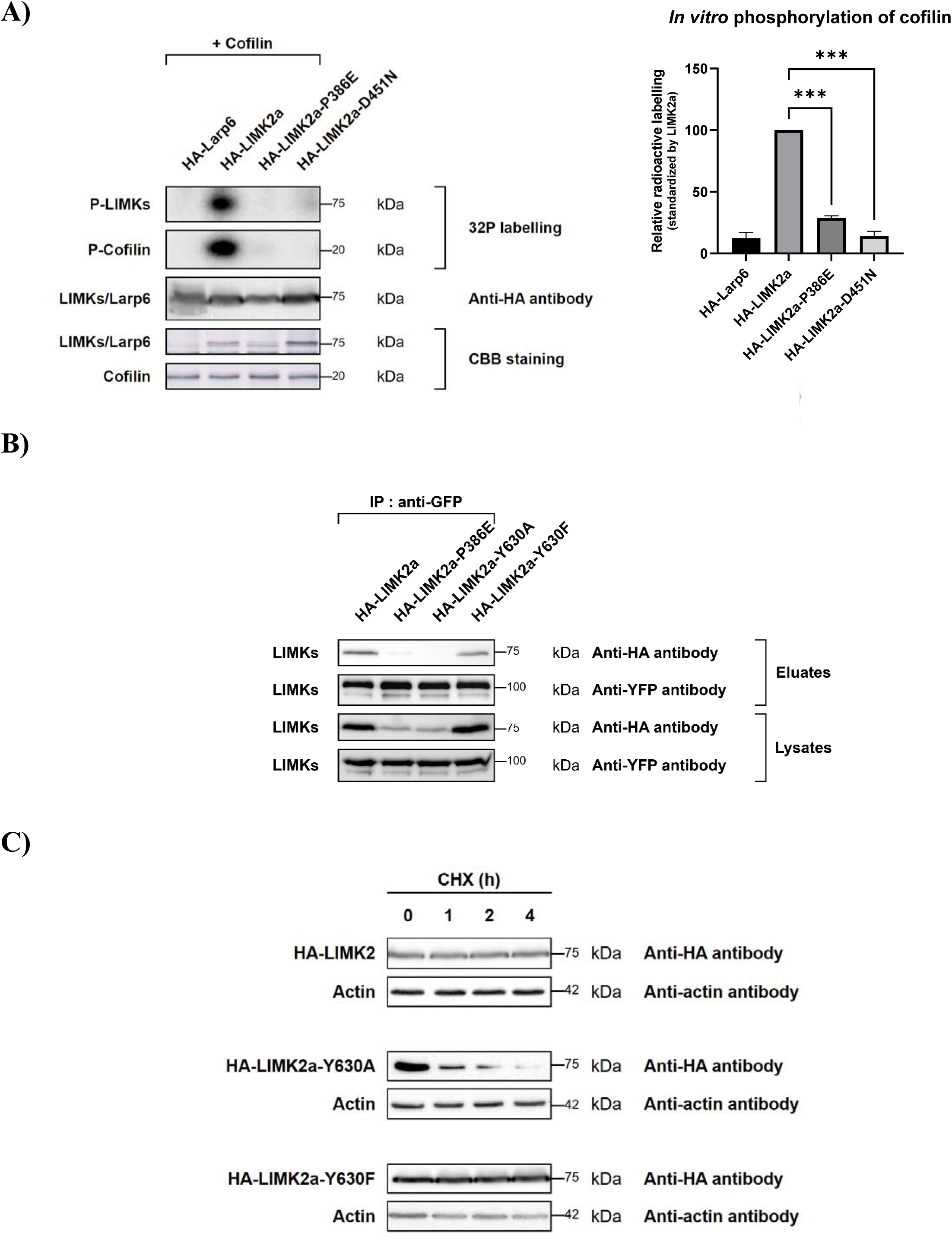
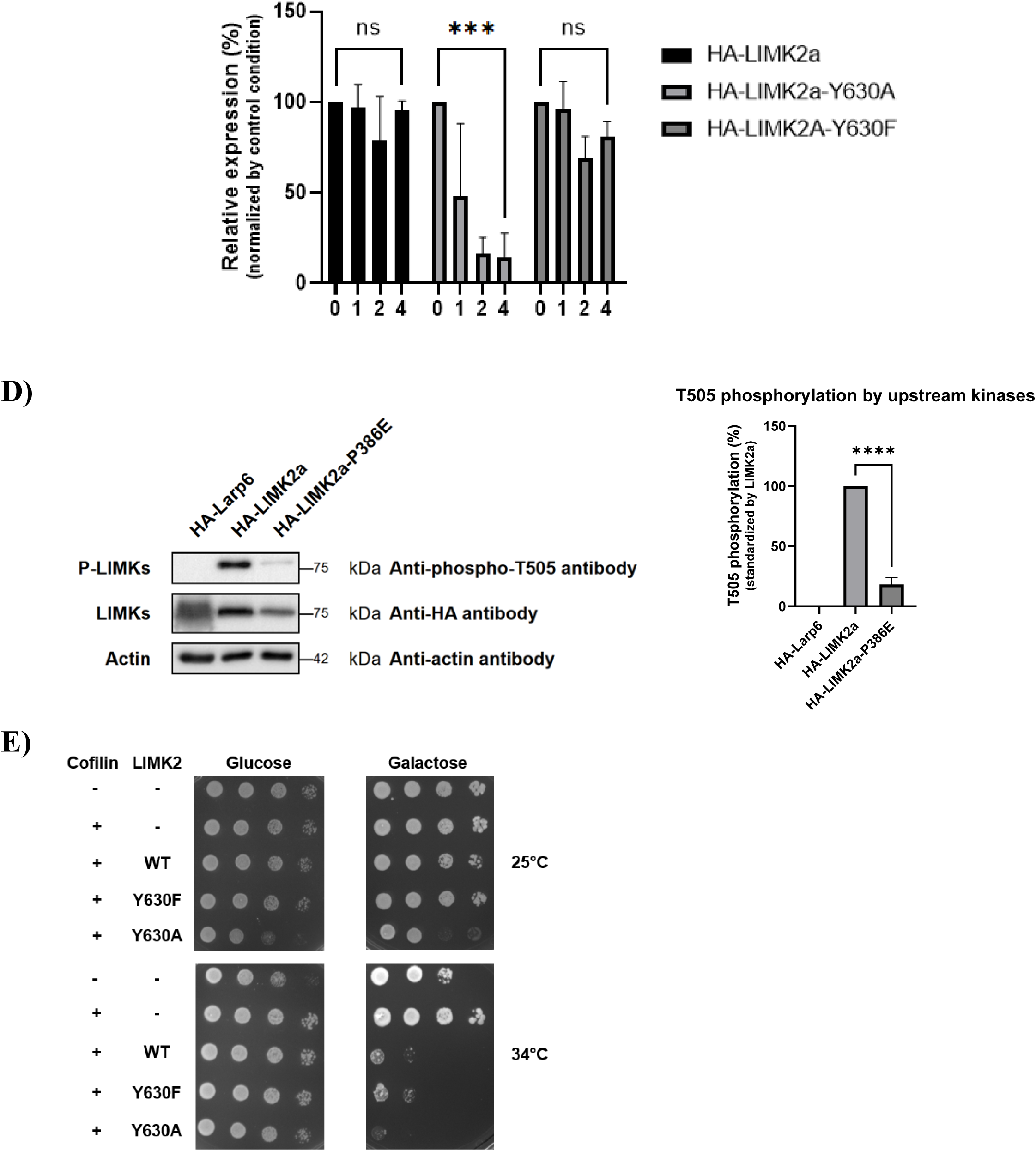
Y630 plays a critical role for LIMK2 dimerization and subsequent transphosphorylation leading to its optimal activity on cofilin. A) *LIMK2 self-phosphorylation occurs by transphosphorylation.* HEK-293 cells overproduced indicated proteins. Anti-HA immunoprecipitated proteins were used in the kinase assay. The anti-HA immunoprecipitates were also subjected to anti-HA immunoblotting and Coomassie blue staining. Quantification of cofilin phosphorylation as depicted in Figure 2B is shown in the right panel, upper graph. Quantification of LIMK2a phosphorylation as depicted in Figure 2D is shown in the right panel, lower graph. B) *Y630 is involved in LIMK2 ability to dimerize.* HEK-293 cells overproduced YFP wild-type LIMK2a and indicated HA-tagged LIMK2a constructs. An immunoprecipitation with anti-GFP grafted beads was performed. Lysates and eluates were subjected to western blotting. C) *LIMK2a-Y630A mutant is less stable than wild type or LIMK2a-Y630F.* Top panel: HEK-293 cells overproduced indicated proteins. Cycloheximide was added at specific time points (4h, 2h, 1h and 0h) before lysis. Lysates were subjected to western blotting. Bottom panel: LIMK expression was quantified. LIMK valuess at time 0 was set at 100%. D) *Transphosphorylation is a prerequisite for T505 phosphorylation.* HEK-293 cellsoverproduced with indicated proteins and cMyc-tagged ROCK. Lysates were subjected to western blotting. Quantification of T505 phosphorylation of LIMK2 as depicted in Figure 4C is shown in the right graph. E) *Yeast complementation assay. Cof^ts^* yeast strain transformed with the indicated expression plasmids were grown on either glucose or galactose to induce LIMK2 expression at 25°C or 34°C. Image representative of n=3 independent experiments.

### Y630 plays a crucial role for LIMK2 dimerization, stability and subsequent transphosphorylation

Subsequently, we wanted to assess if Y630 could play a role in LIMK2a’s ability to dimerize and transphosphorylate similarly to P394 in LIMK1 [69].

First of all, we studied if Y630 was involved in LIMK2a dimerization. We tested the interaction between wild-type LIMK2a and LIMK2a mutants Y630A or Y630F by co-immunoprecipitation experiments. These HA-tagged constructions were co-transfected with YFP-tagged wild-type LIMK2a. An immunoprecipitation with anti-GFP grafted beads was performed. As expected, wild-type LIMK2a interacted together, and LIMK2a-P386E did not interact with LIMK2a. Concerning the Y630 mutants, the LIMK2a-Y630F mutant interacted with wild-type LIMK2a, whereas the LIMK2a-Y630A mutant did not (**Figure 6B**). These results show that Y630 mutation into alanine abolishes LIMK2a ability to homodimerize.

It was previously shown that LIMK1 stability is correlated to its ability to dimerize ^67^. Mutants unable to dimerize were shown to have a very reduced lifespan of approximately 4 hours ^67^ whereas LIMK2a is a very stable protein with a lifespan of around 20 hours ^68^. Therefore, we evaluated the lifespan of LIMK2a-Y630A and LIMK2a-Y630F and compared it to that of wild-type LIMK2a. We performed a cycloheximide chase assay. HEK cells were transfected with one of the different HA-tagged mutants of LIMK2a and incubated with cycloheximide for different times. Cells were lysed and lysates were analysed by western blot with appropriate antibodies. In our experiments, the LIMK2a-Y630A signal rapidly and drastically decreased. Upon 1 hour of cycloheximide treatment, it dropped to 50% and reached 10% upon 2 hours of treatment. (**Figure 6C**). On the contrary, the LIMK2a-Y630F signal was constant all along the time course of the experiment similar to the wild-type LIMK2a signal, suggesting that the LIMK2a-Y630F lifespan was not affected and that both wild-type and LIMK2a-Y630F behave the same way in this range of time (**Figure 6C**).

Altogether, our results showed that LIMK2a-Y630A dimerization is abolished leading to a loss of its ability to transphosphorylate and to phosphorylate cofilin. We also showed that phosphorylation of the canonical T505 is lost in the LIMK2a-Y630A mutant (**Figure 4C**). We then checked this feature for the LIMK2a-P386E mutant by western-blot. This mutant was not phosphorylated on its T505 (**Figure 6E**). These results suggested that T505 phosphorylation is dependent on LIMK2a ability to dimerize and transphosphorylate, Y630 and P386 playing a crucial role in these processes, which appear as indispensable prerequisites for LIMK2a phosphorylation on its T505 and subsequent activation and optimal activity on cofilin.

### Physiological effects of LIMK2a-Y630A/F mutants on a cof^ts^ yeast strain

In order to further explore the physiological role of Y630, we took advantage of a well-characterized *cof1^ts^* yeast strain constructed by D. G. Drubin’s lab ^60^ and adapted an assay previously developed by Boggon’s lab ^69,70^. In the yeast *Saccharomyces cerevisiae*, *COF1* is an essential gene, its deletion is lethal. D. G. Drubin constructed a strain harbouring point mutations in the *yCOF1* gene leading to a thermosensitive strain, *cof1^ts^*. At 25 °C, the *cof1^ts^* strain grows well, whereas at temperatures higher than 30 °C, mutant cofilin protein produced by the *yCOF1* mutated gene is no longer functional and the *cof1^ts^* strain is unable to grow. Human Cofilin can revert this phenotype, as the *cof1^ts^* strain transformed with a plasmid allowing the production of hCofilin is able to grow at high temperatures (above 30 °C). LIMKs are not present in the yeast *S. cerevisiae* as they appeared later in evolution. However, when human LIMK1 is overproduced in yeast, it is able to phosphorylate yeast cofilin on its S3 resulting in its inactivation, which is deleterious for yeast. As a consequence, when LIMK1 is overproduced concomitantly with hCofilin in the *cof1^ts^* strain, it suppresses hCofilin-induced growth restoration and leads to yeast death at temperatures higher than 30 °C. We tested LIMK2a as well as our mutants LIMK2a-Y630A and LIMK2a-Y630F in this system. The *cof1^ts^*strain was transformed with plasmids allowing the constitutive expression of *hCOF1* and the expression of *LIMK2a* wild-type or *Y630A* or *Y630F* mutants under the control of a promoter inducible by galactose. LIMK2 encoded proteins are produced on galactose medium whereas their production is repressed on classical glucose growth medium. We tested the growth ability of each of these strains by performing a drop dilution assay. At 25 °C, on glucose medium, all of the different strains grew as yCofilin is functional. However, we observed a growth defect for the strain co-transformed with plasmids allowing the production of hCofilin and LIMK2a-Y630A (**Figure 6E**). In these conditions (Glucose medium), gene encoding LIMK2a-Y630A should be repressed but it is well known that galactose promoter is a bit leaky and allows a slight production of the protein under its control, even on Glucose medium. This slight production of LIMK2a-Y630A seemed to be detrimental for yeast growth. This phenotype was enhanced on Galactose medium, which induced LIMK2a-Y630A expression. At 34 °C on glucose medium, all the strains grew the same way, even the strain carrying plasmids allowing the production of hCofilin and LIMK2a-Y630A. At 34 °C on galactose medium, the strain transformed with the plasmid allowing only the production of hCofilin still grew efficiently as the hCofilin complements endogenous yeast cofilin defect. On the contrary, the strains co-transformed with plasmids allowing the co-production of hCofilin and of the different mutants of LIMK2a grew poorly. In these conditions, as expected, wildtype LIMK2a phosphorylated cofilin, leading to its inactivation, and severely impaired yeast growth. LIMK2a-Y630F exhibited the same growth profile as wildtype LIMK2a suggesting that this mutant behaves the same way as wild type LIMK2, and is active on cofilin. LIMK2a-Y630A grew the poorest. As this mutant did not phosphorylate cofilin in mammalian cells, we expected that it would not impaired yeast growth in these conditions and behave like the strain only transformed with hCofilin. It seems that the growth defect induced by LIMK2a-Y630A is due to a general deleterious physiological response of yeast when LIMK2a-Y630A is expressed, reinforcing the crucial role of Y630 for LIMK2a function. However, this deleterious effect was not observed at 34 °C on Glucose medium, suggesting that yeast heat-shock response may counterbalance small amounts of LIMK2-Y630A, which might have a folding defect.

### Specific conformations of Y630/Y632 in LIMK2/1

We went further in our study by investigating the impact of Y630 and Y632 respectively on LIMK2a and LIMK1 conformations.

We superimposed all of the 3D structures of the kinase domains of LIMK1 and LIMK2 resolved by X-ray crystallography and deposited in the protein databank (see **Figure 7, upper panel**). It appeared that Y630/Y632 adopt two different well-defined conformations: so-called “in” and “out” conformations depending on the orientation of its side chain towards the core of the protein or towards the solvent (**Figure 7, upper panel**). Even more interestingly, the conformation of Y630/Y632 seems to be correlated to the status of the αC-helix of LIMKs, a conserved structural motif involved, along with the DFG motif, in the balance between active and inactive states of the protein. Indeed, in 13 of these 14 structures when Y630/Y632 adopts an “out” position, the αC-helix adopts an in-conformation, meaning that the conserved glutamic acid (E384 in LIMK1 or E376 in LIMK2) in the αC-helix forms a salt bridge with the conserved b3 lysine (K368 in LIMK1 or K360 in LIMK2). On the contrary, when Y630/Y632 adopt an “in” position, the αC-helix is in an out-conformation, meaning that the glutamic acid of the αC-helix is turned away from the active site, leading to a displacement of the α C-helix. (**Figure 7, lower panel**). This analysis was performed on all chains of the LIM kinase domain of these 14 PDB structures, conducting to observe this trend 32 times from the 33 LIM kinase domain crystal structures. The only structure presenting both αC helix and Y632 in an “in” conformation is the mutated D460N LIMK1 structure of PDB ID 5HVJ. These results suggest that Y630/Y632 and αC-helix conformations may influence each other. As mentioned before, the conformation of residues belonging to the DFG motif and αC-helix mediates the activity status, inactive or active, of kinases, which is also linked to the phosphorylation of the activation loop, corresponding to T505/T508 for LIMK2 and LIMK1, respectively. Our results showed that LIMK2a-Y630A is not phosphorylated on its T505, whereas LIMK2a-Y630F is. These data suggest that the aromatic ring of Y630 may play a crucial role for LIMK2 conformation and activation on its T505.

**Figure 7:**
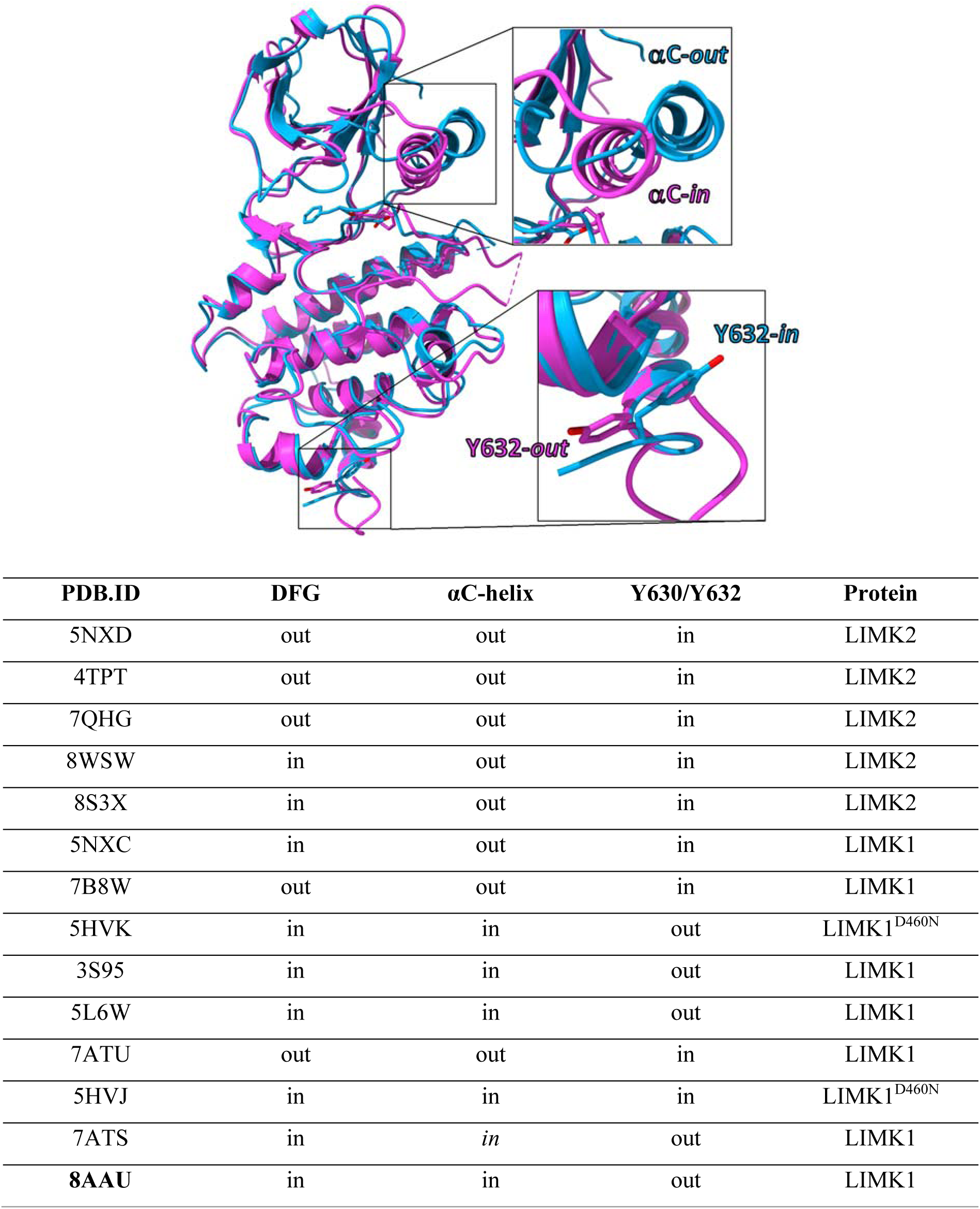
Structural analysis of Y632 and Y630. *Y632/Y630 and αC-helix conformations seem to be correlated in LIMK1/LIMK2 3D-structures.* Upper panel: superposition of two representative 3D structures of LIMK1 kinase domain. The PDB ID 7B8W is represented in blue ribbons and the PDB ID 3S95 in pink ribbons. Orientation of Y632 and αC-helix are zoomed in. Lower panel: Report of the conformations of DFG motif, αC-helix and tyrosine 632/630 for all existing LIMK structures in the RCSB PDB.

To evaluate the stability of the Y630/Y632 “in/out” conformations, we performed molecular dynamics simulations on a representative system of each type of conformation. We chose the LIMK1 structure of PDB ID 3S95 as representative of the Y632-“out”conformation whereas the LIMK1 structure of PDB ID 5NXC represents the Y632-“in” conformation. During the 2 microseconds of simulation, both systems remain globally stable as confirmed by the RMSD and RMSF analyses (see figures S1 and S2 of the supplementary data). The evolution of the Y632 conformation during the simulation is followed by calculating the variation during the simulation of the dihedral angle θ_Y_ defined by the atoms N, C_α_, C_β_ and C_γ_ of the residue (see figure S3 of the supplementary data). The values adopted by this dihedral angle were first evaluated on the Y630/Y632 residues of the crystallographic structures of LIMKs. We observed that the “out” conformation of Y630/Y632 is characterized by θ_Y_=192±9° while the “in” conformation adopts a value of θ_Y_=302±12°. The “in” conformation of Y632 from PDB ID 5NXC remains very stable during the simulation, with a mean value of the dihedral angle of θ_Y_=296±13°. Little more variations are observed for the simulation of the Y632-out conformation from PDB ID 3s95 with very transitory switches between conformations “out” and “in”, leading to a mean value of the dihedral angle of θ_Y_=196±46°. Thus, these molecular dynamics simulations clearly demonstrated the persistence of these two conformations for residue Y630/632 which are therefore not an artifact of crystallization, what reinforces our conviction of a structural role of Y630/Y632.

## Discussion

The LIM kinases, LIMK1 and LIMK2, play a major role in cytoskeleton dynamics especially by regulating the turnover of actin filaments via the phosphorylation and subsequent inhibition of cofilin, an actin depolymerizing factor, their canonical substrate in this pathway. Consequently, LIM kinases are involved in many physiological processes as well as several pathologies ^54^. Although LIM kinases have appeared as major therapeutic targets, they remain undruggable as no small molecules inhibiting their kinase activity have successfully gone through clinical trials so far ^48^. It is thus vital to have a better understanding of the activity and regulation of these proteins to develop successful new therapeutic approaches and further develop our knowledge of these proteins.

In our current study, we have pointed out a new regulation mechanism of for LIM kinases. A single residue, Y630 for LIMK2 and Y632 for LIMK1, appeared to play a crucial role for the optimal activity of these proteins on cofilin. Y630/Y632 mutation into an alanine led to a loss of kinase activity of LIMKs on cofilin, along with a loss of phosphorylation of the full-length protein. However, the kinase activity of these mutants was preserved on the broad kinase substrate Myelin basic Protein (MBP), which suggests that this process of regulation is specific to the substrate cofilin. A loss of the propensity of LIMKs to induce actin stress fiber formation was also observed with these mutants. Moreover, Y630/Y632 are well conserved within LIMK sequences throughout evolution, which is consistent with a potential vital role of these amino acids. The crucial role of Y630/Y632 was also supported by experiments on yeast. Indeed, the Y630A mutant appeared to have deleterious effects on yeast growth under normal growth conditions at 25 °C. This growth defect was no longer observed at higher temperatures (34 °C) suggesting that the stress response machinery may counteract the deleterious effects observed with Y630A. When the Y630A mutant was overexpressed, a growth defect was also observed at both temperatures reinforcing the adverse impact of this mutation on LIMK function.

We initially thought that Y630/Y632 were potential phosphorylation sites. Indeed, when they were mutated into an alanine, phosphorylation of full-length LIMKs was no longer observed in our *in vitro* assay. Early studies on LIMKs have pointed out a global phosphorylation of LIMK1 mainly on serine and tyrosine ^71^. When we mutated Y630 into a phosphorylatable (S) or phosphomimetic amino acid (E) we did not restore either cofilin phosphorylation or full-length LIMK phosphorylation. We then hypothesized that these tyrosines may rather play a structural role due to their aromatic ring. Indeed, when Y630/Y632 were mutated into a phenylalanine, these mutants behaved the same way as the wild-type LIMKs: they phosphorylated cofilin, and were phosphorylated on their own. By superimposing the 3D structures of LIMK1 and LIMK2 available in the PDB, it appeared that Y630/Y632 adopt two discrete conformations we named “in” and “out” according to the orientation of its aromatic ring towards the core of the protein. This conformation seemed to be correlated to the one of the α-C-helix, which also adopts two discrete conformations “in” and “out” and to a lesser extent to the DFG “in” and “out” conformations (Figure 7). α-C-helix conformation from the N-lobe of kinases is known to play a crucial role in their activity status, the “in” conformation corresponding to the active form of the kinase ^72^. Furthermore, an α-C-helix is known to mediate dimerization as described for EGFR and B-Raf ^73–75^. We showed that Y630/Y632 play a crucial role for LIMK dimerization, LIMK2-Y630A being unable to dimerize whereas LIMK2-Y630F did dimerize. This dimerization could therefore be mediated by the α-C-helix whose conformation would be conditioned by the one of Y630/Y632. LIMKs were shown to sometimes dimerize via the mediation of other partners such as Hsp90 [66] or TrkB ^76^. Here, we cannot rule out LIMK dimerization mediated or not by another partner, as we were working on overexpressed proteins in mammalian cells. Li et al. ^67^ showed that this dimerization had an impact on LIMK lifespan and phosphorylation. Concordantly, we have shown that LIMK2-Y630A exhibited a very short lifespan compared to LIMK2 wild type or LIMK2-Y630F. Taking advantage of different mutants of LIMKs described in the literature and previously characterized, we demonstrated that LIMK dimerization resulted in its transphosphorylation. Similarly, former studies have shown that LIMK1 phosphorylation was strongly decreased in its kinase dead version (D460A), phospho-Ser and phospho-Tyr sites being mainly affected ^71^. We tried to identify these phosphorylation sites, focusing on previously known phosphorylation sites described in the literature, S283 and T494 for LIMK2 [73]. We constructed the double mutant LIMK2-S283A-T494A, which still phosphorylated cofilin in our *in vitro* assay, as well as the triple mutant LIMK2-S283A-T494A-T505A, which was also was still phosphorylated in our *in vitro* assay, suggesting another site or sites of possible phosphorylation existing in LIMK2. Furthermore, we showed that LIMK2-T505EE constitutive activity was lost in the presence of the extra mutation Y630A, LIMK2-Y630A-T505EE being unable to phosphorylate cofilin (**figure 4D**). Altogether, our results suggest that Y630/Y632 mediate transphosphorylation of LIMK on other residues than the canonical threonine belonging to the activation loop. This (these) phosphorylation(s) would be a prerequisite for subsequent T505/T508 phosphorylation leading to optimal activity of LIMKs towards cofilin as LIMK2-T505EE-Y630A is no longer able to bypass T505 activation contrary to LIMK2-T505EE. These data point out a new mechanism of regulation of the kinase activity of LIMKs specifically on cofilin.

LIMK activation was initially alleviated to the phosphorylation of their threonine T505/T508 belonging to their activation loop ^9,53,71,77^. However, a growing body of evidence suggests that the regulation of these kinases is much more complex, and LIMKs appear to have unusual characteristics compared to other kinases.

LIMKs possess an atypical consensus kinase sequence which made them impossible to classify as Ser/Thr kinases or Tyr kinases when they were first discovered (DLNSHN sequence for LIMKs vs DLKxxN sequence for Ser/Thr kinases and DLRAAN/DLAARN sequence for Tyr kinases) ^50,78^. Later on, it was shown that LIMKs were able to phosphorylate both serine (as shown for cofilin, SRPK1, PTEN, SPOP, TWIST1, NKX3.1) and tyrosine (as shown for MT1-MMP and mutated cofilin-S3Y) ^54,79^. Furthermore, their active site is quite shallow as an asparagine replaces the usual lysine present at the HRD+2 position in the catalytic loop, accommodation of the substrate is very tight ^80^. However, it was recently shown based on LIMK1/cofilin complex structure that a rock-and-poke mechanism allows different position of the phospho-acceptor residue into the active site of LIMKs explaining the dual kinase activity on both serine and tyrosine ^79^. In early studies, an allosteric regulation of LIMK kinase activity by their LIM domains was described. LIMK1 LIM domain was shown to interact with its kinase domain ^81^, leading to its inhibition ^82^. Indeed, a LIM-truncated mutant of LIMK1 much more phosphorylated efficiently cofilin compared to wild-type LIMK1, and had a drastic effect on actin cytoskeleton organization. Furthermore, when the cysteine of the zinc fingers of the LIM domains of LIMK1 were mutated into glycine leading to the loss of LIMK zinc finger tertiary structure, a drastic increase of cofilin phosphorylation was observed. These data were further confirmed by the studies of Edwards et al. ^77^ who showed that deletion of the LIM domains had a strong impact on actin cytoskeleton remodelling. In this study, authors also showed that the PDZ domain of LIMK1 inhibited its kinase activity as a GL177EA mutant led to actin aggregation and increased cofilin phosphorylation. This mutant was designed by homology with the third PDZ domain of PSD-95, which interacts with the C-terminus of the potassium channel via its GLGF sequence. The 3D structure of LIMK2 PDZ domain has recently been resolved by X-ray crystallography, thus corroborating this data ^70^. PDZ domains are common structural features found in many proteins. Deprived of any catalytic activity, they mediate protein/protein interactions via their binding site forming a groove between an α-B-helix and a β-B-strand, where a conserved motif x-ϕ-G-ϕ‘ (x being any residue, and ϕ an hydrophobic residue) plays a crucial role in partner recognition. Actually, the G residue of the x-ϕ-G-ϕ‘ sequence corresponds to G177 mutated in Edwards et al’s studies having a notable effect on LIMK activity underlying the crucial role of this sequence. Intriguingly, LIMKs partially possess this sequence as they are the sole proteins for which the first hydrophobic residue ϕ is replaced by an Arg. This Arg retains the usual ϕ orientation towards the hydrophobic core of the protein, the presence of its charge is counterbalanced by an extensive hydrogen network leading to a rigidity of the α-B-helix and consequently to a shallow groove limiting LIMK interaction with partners. Furthermore, an extended distal conserved surface was also identified to play a major role in LIMK activity regulation as mutation of residues of this area resulted in an increased phosphorylation of cofilin by these mutants. This structure revealed intriguing features showing that the LIMK PDZ domain differs from canonical PDZ domains in several points and contributes to LIMK negative autoregulation. Recently, the group of Boggon pointed out another feature of LIMKs negatively regulating their activity ^83^. They showed that the presence of T508 in the activation loop of LIMK1 also lowers its activity. Indeed, when T508 was mutated into a serine, PAK4 phosphorylated this residue much more efficiently. T508 contributes to the sub-optimal activation of LIMK1 by PAK4. Indeed, this feature was already observed in initial studies on LIMK1, where Ohashi et al. ^71^ observed an increased phosphorylation of the mutant LIMK1-T508S compared to its wild-type version. Another intriguing feature about LIMKs is their atypical interaction with their canonical substrate cofilin allowing a high-fidelity exclusive kinase-substrate recognition ^69,79^. As LIMKs phosphorylate cofilin on its Ser3, it is not possible for them to recognize their target thanks to amino acids flanking it as it is usually the case for many kinase-substrate couples. They bypass this usual required mechanism by an atypical recognition mechanism: a very selective and specific complementary interaction of the Ser3-sequentially-distal α5-helix of cofilin with a groove of LIMK1 leads to the tilting of cofilin-Ser3 within LIMK1 active site leading to its phosphorylation. This fine-tuned mechanism of kinase-substrate recognition had never been observed prior to these two complementary and consistent studies.

It appears that LIMK activity is finely tuned by many different unusual molecular features reflecting a strong ability of life to precisely control every processes. Most of the mechanisms of LIMK regulation so-far have led to an inhibition or drastic decrease of LIMK kinase activity towards cofilin. Here, we pointed out an activation mechanism absolutely required for LIMK kinase activity towards cofilin. These molecular features bring new insights into LIMK regulation and activity understanding.

## Author Contributions

Original draft preparation and writing, E.V. and B.V.; All authors have read and agreed to the published version of the manuscript.

## Acknowledgments

We would also like to thank the RTR Motivhealth, the Ligue contre le Cancer Grand Ouest (comités d’Eure et Loire et du Loiret), Cancéropole Grand Ouest, Réseau Molécules marines, métabolisme et cancer for their support. Authors thanks the projects CHemBio (FEDER-FSE 2014-2020-EX003677), Valbiocosm (FEDER-FSE 2014-2020-EX003202), Techsab (FEDER-FSE 2014-2020-EX011313), QUALICHIM (APR-IA-PF 2021-00149467), Project ESTIM-ICOA (CPER / FEDER-FSE+ 2021-2027-00022860), the RTR Motivhealth (2019-00131403) and the Labex programs SYNORG (ANR-11-LABX-0029) and IRON (ANR-11-LABX-0018-01) for their financial support of ICOA, UMR 7311, University of Orléans, CNRS. Figures were created with Sciugo.com. Many thanks to Karen Plé for her English proofreading.

