## Supplementary material for "A new mechanism of regulation of LIM kinases, LIMK1 and LIMK2, modulates their activity on cofilin and actin filament remodelling": Sup Data

### Supplementary data

| Plasmid | Description | Reference |
|---|---|---|
| **pcDNA3-(HA)2-LARP6** | PCMV-(HA)2-LARP6 | Vallée et al., 2018 |
| **pcDNA3-(HA)2-LIMK2-2a** | PCMV-(HA)2-LIMK2-2a | Vallée et al., 2012 |
| **pcDNA3-(HA)2-LIMK2-2a-ΔCT591** | PCMV-(HA)2-LIMK2ΔCT591 | This work |
| **pcDNA3-(HA)2-LIMK2-2a-ΔCT602** | PCMV-(HA)2-LIMK2ΔCT602 | This work |
| **pcDNA3-(HA)2-LIMK2-2a-ΔCT609** | PCMV-(HA)2-LIMK2ΔCT609 | This work |
| **pcDNA3-(HA)2-LIMK2-2a-ΔCT611** | PCMV-(HA)2-LIMK2ΔCT611 | This work |
| **pcDNA3-(HA)2-LIMK2-2a-T625A** | PCMV-(HA)2-LIMK2-2a-T625A | This work |
| **pcDNA3-(HA)2-LIMK2-2a-S627A** | PCMV-(HA)2-LIMK2-2a-S627A | This work |
| **pcDNA3-(HA)2-LIMK2-2a-Y630A** | PCMV-(HA)2-LIMK2-2a-Y630A | This work |
| **pcDNA3-(HA)2-LIMK2-2a-T633A** | PCMV-(HA)2-LIMK2-2a-T633A | This work |
| **pcDNA3-(HA)2-LIMK2-2a-S636A** | PCMV-(HA)2-LIMK2-2a-S636A | This work |
| **pEN1-(YFP)** | pCMV-(YFP) | Vallée et al., 2018 |
| **pEN1-(YFP)-LIMK2-2a** | pCMV-(YFP) | Vallée et al., 2018 |
| **pEN1-(YFP)-LIMK2-2a-Y630A** | pCMV-(YFP) | This work |
| **pcDNA3-(HA)2-LIMK1** | pcDNA3-(HA)2-LIMK1 | Vallée et al., 2012 |
| **pcDNA3-(HA)2-LIMK1-Y632A** | pcDNA3-(HA)2-LIMK1-Y632A | This work |
| **pcDNA3-(HA)2-LIMK2-2a-Y630S** | PCMV-(HA)2-LIMK2-2a-Y630S | This work |
| **pcDNA3-(HA)2-LIMK2-2a-Y630E** | PCMV-(HA)2-LIMK2-2a-Y630E | This work |
| **pcDNA3-(HA)2-LIMK2-2a-Y630F** | PCMV-(HA)2-LIMK2-2a-Y630F | This work |
| **pcDNA3-(HA)2-LIMK2-2a-T505A** | PCMV-(HA)2-LIMK2-2a-T505A | This work |
| **pcDNA3-(HA)2-LIMK2-2a-T505EE** | PCMV-(HA)2-LIMK2-2a-T505EE | This work |
| **pcDNA3-(HA)2-LIMK2-2a-P386E** | PCMV-(HA)2-LIMK2-2a-P386E | This work |
| **pcDNA3-(HA)2-LIMK2-2a-D451N** | PCMV-(HA)2-LIMK2-2a-D451N | This work |
| **pcDNA3-(HA)2-LIMK2-2a-P386E-Y630F** | PCMV-(HA)2-LIMK2-2a-P386E-Y630F | This work |
| **pcDNA3-(HA)2-LIMK2-2a-T505A-Y630F** | PCMV-(HA)2-LIMK2-2a-T505A-Y630F | This work |
| **pRS426-COF1** | PGAL1-COF1 | This work |
| **pRS426- COF1-LIMK2** | PGAL1-COF1-LIMK2-2a | This work |
| **pRS426- COF1-LIMK2-Y630A** | PGAL1-COF1-LIMK2-2a-Y630A | This work |
| **pRS426- COF1-LIMK2-Y630F** | PGAL1-COF1-LIMK2-2a-Y630F | This work |

**Table 1: Details of the plasmids used in this study**

| **Construction** | **Origin vector** | **Primer** | **Sequence (5'-3')** | **Fragment lenght** |
|---|---|---|---|---|
| **HA-LIMK2-2a-Y630A** | HA-LIMK2-2a | For | GAGCATGCAGGCCGGCCTGACC | 7286 |
| | | Rev | ACAGTGTGGTCCAACTCC | |
| **YFP-LIMK2-2a-Y630A** | HA-LIMK2-2a-Y630A | For | CGACGTGAATTCGCCACCATGTCCGCGCTGGCGGGTGAA | 1947 |
| | | Rev | CTGAGCCCGCGGTGAGGGAGGTGAGTCCCGGGTCA | |
| **HA-LIMK1-Y632A** | HA-LIMK1 | For | CTGGGAGACCGCCCGGCGCGGC | 7385 |
| | | Rev | AAACCTCTGTCCAGCTGCTC | |
| **HA-LIMK2-2a-T505A** | HA-LIMK2-2a | For | GAAGCGCTACGCGGTGGTGGG | 7286 |
| | | Rev | TTGCGGTCGTTCTTGCGC | |
| | | For | GAGCATGCAGTCCGGCCTGAC | 7286 |

| | | | | |
|---|---|---|---|---|
| **HA-LIMK2-2a-Y630S** | HA-LIMK2-2a | Rev | ACAGTGTGGTCCAACTCC | |
| **HA-LIMK2-2a-Y630E** | HA-LIMK2-2a | For | GAGCATGCAGGAGGGCCTGACCC | 7286 |
| | | Rev | ACAGTGTGGTCCAACTCC | |
| **HA-LIMK2-2a-Y630F** | HA-LIMK2-2a | For | GAGCATGCAGTTCGGCCTGAC | 7286 |
| | | Rev | ACAGTGTGGTCCAACTCC | |
| **HA-LIMK2-2a-T505EE** | HA-LIMK2-2a | For | GAAGCGCTACGAGGAGGTGGTGGGAAACCC | 7289 |
| | | Rev | TTGCGGTCGTTCTTGCGC | |
| **HA-LIMK2-2a-P386E** | HA-LIMK2-2a | For | CCTGGACCACGAGAATGTGCTCAAG | 7286 |
| | | Rev | CTGCGCATCACTTTCACC | |
| **HA-LIMK2-2a-D451N** | HA-LIMK2-2a | For | CATCCACCGGAATCTGAACTC | 7286 |
| | | Rev | ATGCACATAGAGTGCAAATAG | |
| **HA-LIMK2-2a-P386E-Y630F** | HA-LIMK2-2a-Y630F | For | CCTGGACCACGAGAATGTGCTCAAG | 7286 |
| | | Rev | CTGCGCATCACTTTCACC | |
| **HA-LIMK2-2a-S283A** | HA-LIMK2-2a | For | GAGGAGACGTGCCCTAAGGCG | 7286 |
| **HA-LIMK2-2a-T494A** | HA-LIMK2-2a | For | CAAGAAACGCGCCTTGCGCAA | 7286 |
| | | Rev | GTGGTGGCCTTCTCCATG | |
| **HA-LIMK2-2a-S283A-T494A** | HA-LIMK2-2a-S283A | For | CAAGAAACGCGCCTTGCGCAA | 7286 |
| | | Rev | GTGGTGGCCTTCTCCATG | |
| **HA-LIMK2-2a-S283A-T494A-T505A** | HA-LIMK2-2a-S283A-T494A | For | GAAGCGCTACGCGGTGGTGGG | 7286 |
| | | Rev | TTGCGGTCGTTCTTGCGC | |
| **COF1** | pRS426-COF1 | For | ATGGCCTCCGGTGTGGCT | 6975 |
| | | Rev | TCACAAAGGCTTGCCCTCCAGGG | 6975 |
| **COF1-LIMK2a** | pRS426-COF1-LIMK2 | For | ATGTCCGCGCTGGCGGGT | 9544 |
| | | Rev | CTAGGGAGGTGAGTCCCGGGT | 9544 |

**Table 2: Details of the primers and vectors used in this study**

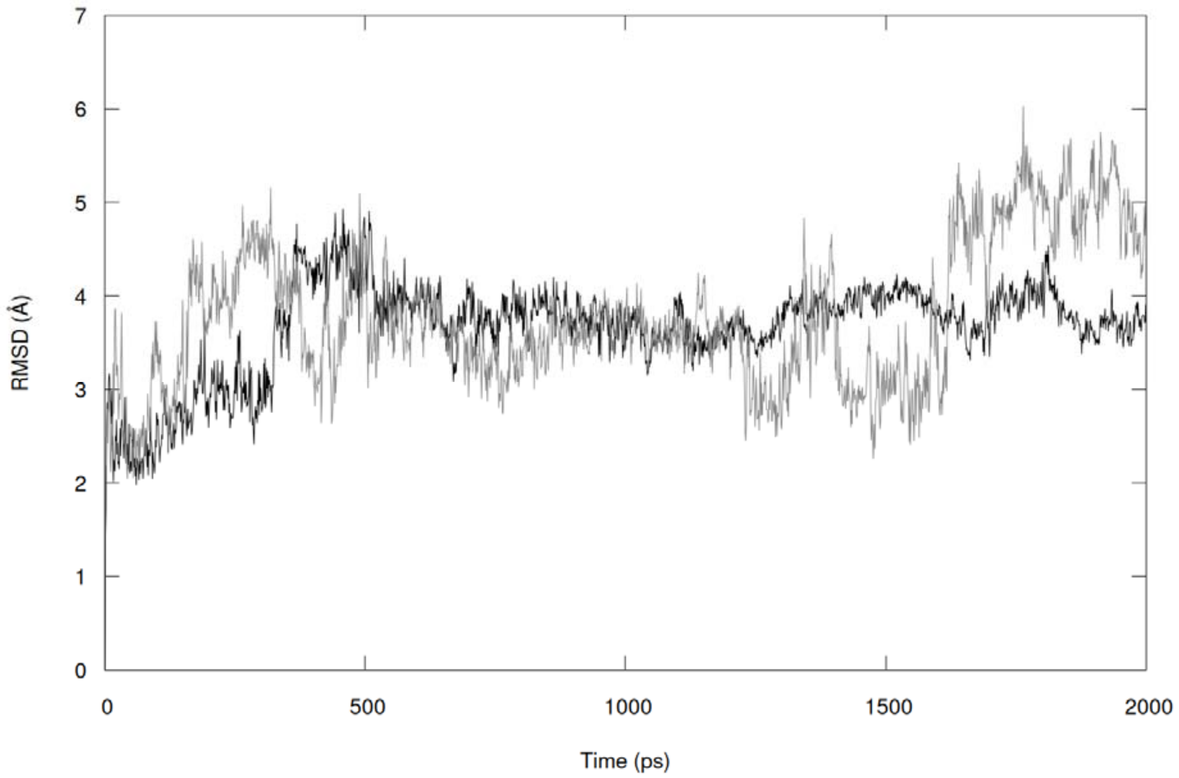


Figure S1: Root mean square deviation (RMSD) calculated on the $C_{\alpha}$ backbone of the Y632-in conformation representative system (black) and of the Y632-out conformation representative system (grey) during 2μs of MD simulation.

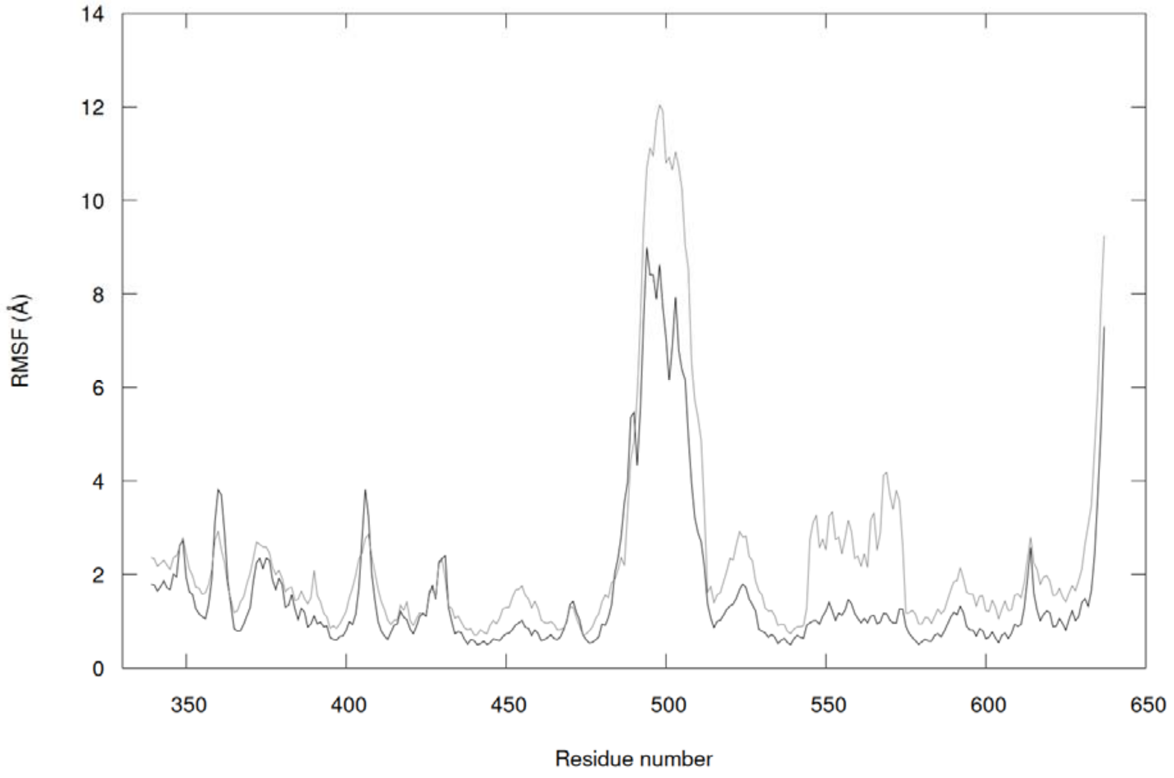

Figure S2: Root mean square fluctuation (RMSF) per residue of the Y632-in conformation representative system (black) and of the Y632-out conformation representative system (grey) during 2μs of MD simulation.

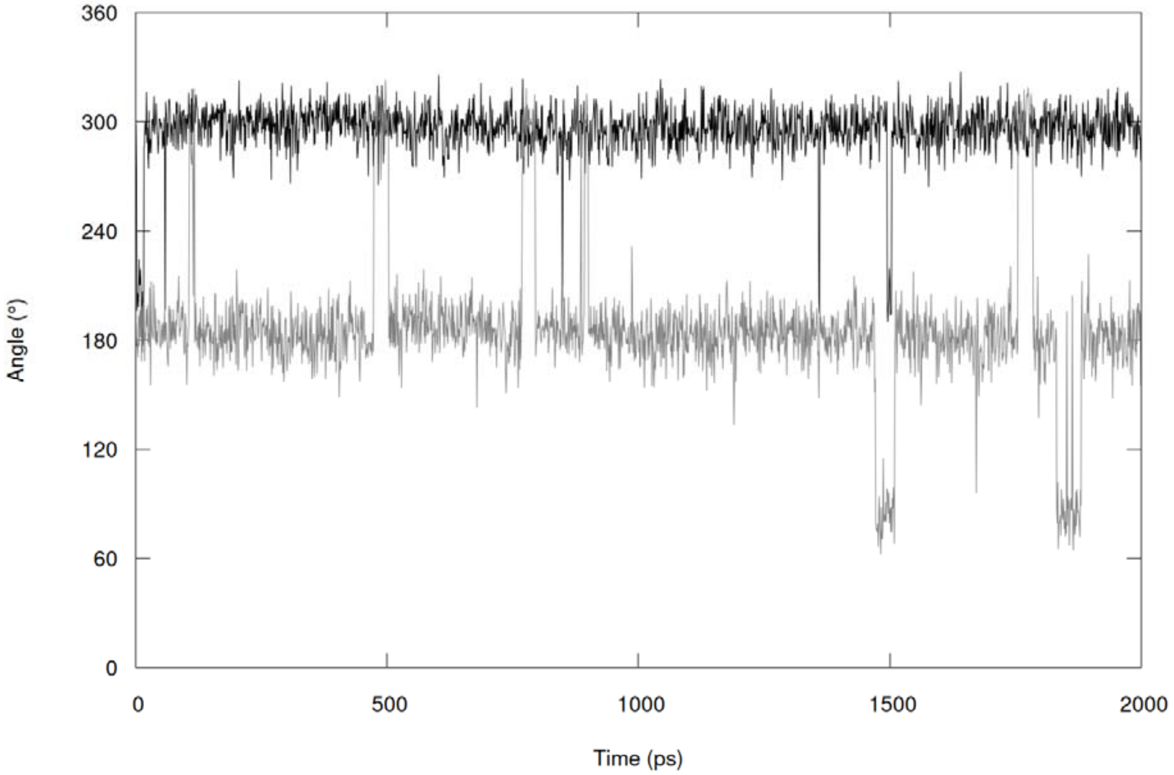


Figure S3: Evolution of the dihedral angle defined by the atoms N, $C_\alpha$, $C_\beta$ and $C_\gamma$ of Y632 for the Y632-in conformation representative system (black) and the Y632-out conformation representative system (grey) during 2μs of MD simulation.
